# Functional redundancy revealed by the deletion of the mimivirus GMC-oxidoreductase genes

**DOI:** 10.1101/2023.04.28.538727

**Authors:** Jean-Marie Alempic, Hugo Bisio, Alejandro Villalta, Sébastien Santini, Audrey Lartigue, Alain Schmitt, Claire Bugnot, Anna Notaro, Lucid Belmudes, Annie Adrait, Olivier Poirot, Denis Ptchelkine, Cristina De Castro, Yohann Couté, Chantal Abergel

## Abstract

The mimivirus 1.2Mb genome was shown to be organized into a nucleocapsid-like genomic fiber encased in the nucleoid compartment inside the icosahedral capsid (1). The genomic fiber protein shell is composed of a mixture of two GMC-oxidoreductase paralogs, one of them being the main component of the glycosylated layer of fibrils at the surface of the virion (2). In this study, we determined the effect of the deletion of each of the corresponding genes on the genomic fiber and the layer of surface fibrils. First, we deleted the GMC-oxidoreductase the most abundant in the genomic fiber, and determined its structure and composition in the mutant. As expected, it was composed of the second GMC-oxidoreductase and contained 5- and 6-start helices similar to the wild-type fiber. This result led us to propose a model explaining their coexistence. Then, we deleted the GMC-oxidoreductase the most abundant in the layer of fibrils to analyze its protein composition in the mutant. Second, we showed that the fitness of single mutants and the double mutant were not decreased compared to the wild-type viruses in laboratory conditions. Third, we determined that deleting the GMC-oxidoreductase genes did not impact the glycosylation or the glycan composition of the layer of surface fibrils, despite modifying their protein composition. Since the glycosylation machinery and glycan composition of members of different clades are different (3, 4), we expanded the analysis of the protein composition of the layer of fibrils to members of the B and C clades and showed that it was different among the three clades and even among isolates within the same clade. Taken together, the results obtained on two distinct central processes (genome packaging and virion coating) illustrate an unexpected functional redundancy in members of the family *Mimiviridae*, suggesting this may be the major evolutionary force behind their giant genomes.

**One-Sentence Summary:** Functional redundancy preserves mimivirus genomic fiber and layer of fibrils formation.

## Introduction

Mimivirus is the inaugural member of the family *Mimiviridae* part of the *Nucleocytoviricota* phylum encompassing large and giant DNA viruses infecting eukaryotes (5). Members of the family *Mimiviridae* infecting amoeba have dsDNA genomes up to 1.5 Mb encoding over 1000 proteins, including a complete glycosylation machinery (3, 4, 6–9). Mimivirus virion penetrate the cell through phagocytosis, and the acidic vacuole mediates opening of the stargate structure at one vertex of its icosahedral capsid (10, 11). The internal membrane unwraps and fuses with the phagosome membrane, allowing transfer of the nucleoid compartment into the host cytoplasm, while empty capsids remain in the vacuole (6, 10, 12).

The infectious cycle occurs in the cytoplasm where a large viral factory is developed (6, 12–14). At the late stage of the cycle, neo-synthesized virions bud at the periphery of the viral factory where they are filled with the genome. Finally, a glycosylated layer of fibrils composed of proteins and two large polysaccharides synthesized by the virally-encoded machinery, is added to the capsids (4, 15, 16). As a result, the 750 nm-diameter virions resemble Russian dolls made of the external layer of reticulated glycosylated fibrils (referred to as the “layer of fibrils”) decorating the surface of the icosahedral capsids. Underneath the capsid shell, the nucleoid compartment encases the 1.2 Mb dsDNA genome organized into a 30 nm large nucleocapsid-like structure (referred to as the “genomic fiber”). The genomic fiber is made of a protein shell internally lined by the folded DNA and a central channel that can accommodate large proteins such as the viral RNA polymerase (1). Three independent genomic fiber structures have been determined by cryo-electron microscopy (cryo-EM): two compact 5- and 6-start DNA-containing helices, and a 5-start relaxed helix, without DNA (1). Unexpectedly, the protein shell was found to be composed of two glucose–methanol–choline (GMC) oxidoreductases sharing 69% sequence identity, with a ratio of 5 between qu_946 and qu_143 according to the protein composition of the purified genomic fiber (1). The resolution of the reconstructions (3.7 and 4.4 Å) prevented us from determining whether each helix contained a single paralog or a mixture of both. Interestingly, one of the two GMC-oxidoreductases, qu_143 (R135 in mimivirus prototype), was known to compose the external layer of fibrils at the surface of mimivirus capsids and it was hypothesized that it was the major target for glycosylation (4, 9, 17, 18). However, despite its involvement in both the mimivirus genomic fiber inside the nucleoid, and the layer of fibrils at the periphery of the icosahedral capsid, GMC-oxidoreductase homologs are absent in the laboratory-evolved mimivirus M4 strain (18). M4 also lacks the glycosylation machinery, described for members of different clades of the subfamily *Megamimivirinae* and responsible for synthesizing and branching the polysaccharides on the capsids (3, 4). In the present study, we used in-house developed tools (19) to delete mimivirus GMC-oxidoreductase genes. We then assessed the fitness cost associated to these deletions and investigated their impact on the formation of the genomic fiber and the protein and glycan composition of the layer of fibrils. Cryo-EM was used to determine the structure of the KO_946 genomic fiber made of qu_143, the less abundant in the wild-type (wt) genomic fiber. Nuclear magnetic resonance (NMR) and Gas Chromatography Mass Spectrometry (GC-MS) were used to analyze the compositions in glycans and their structures for each mutant and to compare them with the wt layer of fibrils. We used Mass spectrometry (MS)-based proteomics to analyze for each of the three mutants the protein composition of their layer of fibrils and extended the study to members belonging to B and C clades (moumouviruses and megaviruses), known to glycosylate their layer of fibrils with different glycans using a clade-specific glycosylation machineries (3, 9). While confirming the non-essentiality of the two GMC-oxidoreductases, our results document the unexpected resilience of mimivirus to the deletion of these two genes through the use of alternative proteins to compensate their disruptions.

## Results

### None of the two GMC-oxidoreductases is essential

We used our recently developed protocol (19) combining homologous recombination with the introduction of a nourseothricin N-acetyl transferase (NAT) selection cassette to delete each of the two genes encoding the GMC oxidoreductases (qu_946 and qu_143). We selected recombinant viruses that were cloned to obtain homogeneous populations (Fig 1B) (19). Each mutant was easily produced and genotyped to confirm the mutation (Fig. S1). Using a second Neomycin resistance gene (NEO) selection cassette we were able to delete both genes (Fig. S1) demonstrating that the two GMC-oxidoreductases were not essential. The absence of additional mutations in every mutant was confirmed by genome sequencing.

**Figure 1:**
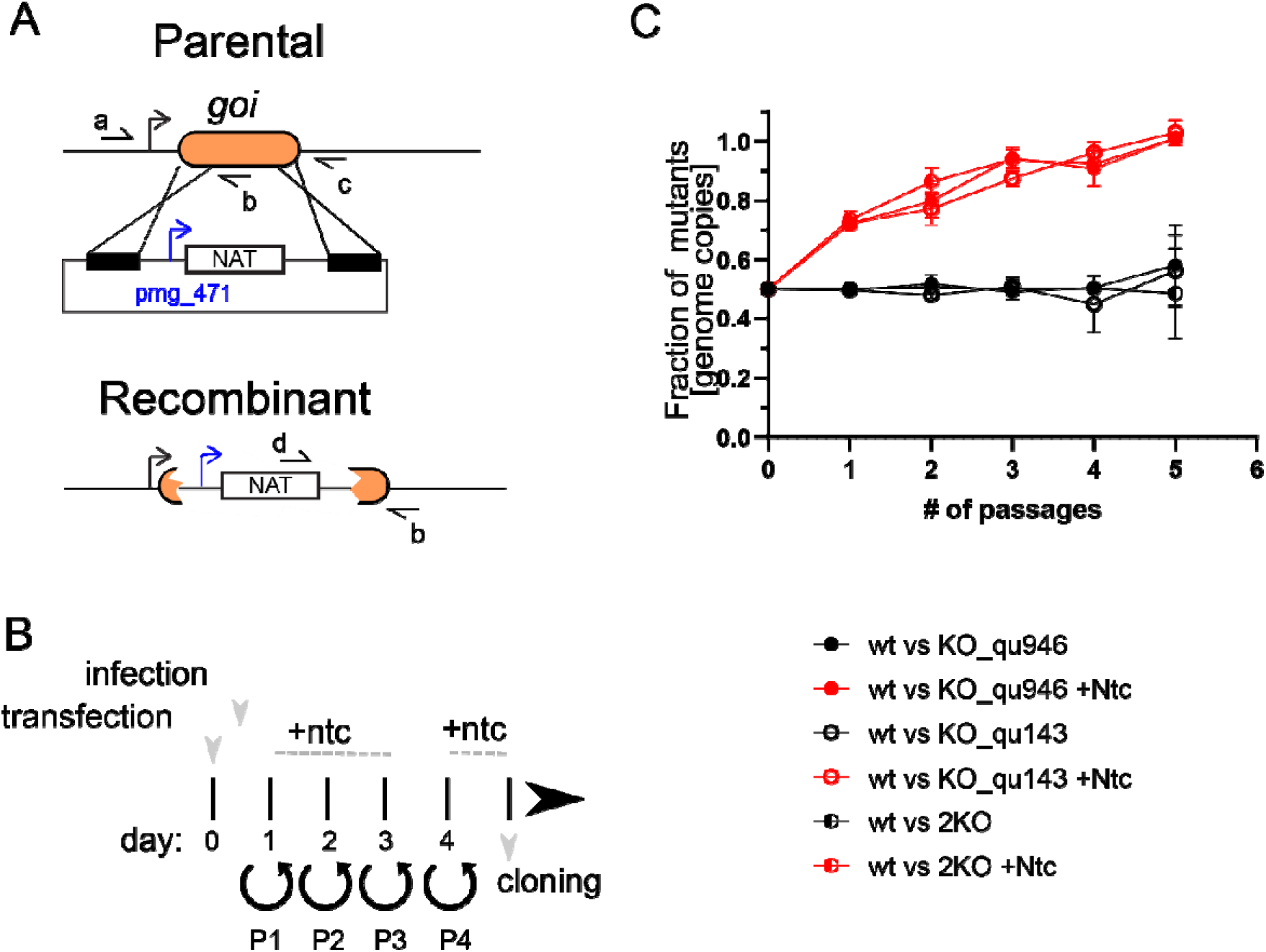
Mimivirus mutants’ generation and phenotypic characterization. A] Schematic representation of the vector and strategy utilized for deletion of qu_143 (KO_qu143) and qu_946 (KO_qu946) in mimivirus reunion strain. *goi*: gene of interest. Selection cassette was introduced by homologous recombination and recombinant viruses were generated as shown in Fig 1B. Primer annealing sites are also shown and the sequence of the primers is included in Table S1. B] Cartoon depicting the strategy for the selection of recombinant viruses. Viral infection was performed 1-hour post-transfection. Ntc: Nourseothricin. P= passage. C] Growth competition assays revealed no significant defects in the lytic cycle of deletion strains. The competition was also performed in presence of Nourseothricin which allows the out competition of the recombinant strains due to the expression of a Nourseothricin selection cassette. Measurements were performed by qPCR of an endogenous locus (present in wt and recombinant strains) and the Nourseothricin selection cassette (only present in recombinant viruses).

### Mutants’ fitness

To assess whether a fitness cost was associated with the mutations, we performed competition assays against wt mimivirus reunion strain by measuring the abundance of each mutant over several cycles in the presence and absence of selection. In contrast to the wt, each mutant presents a common nourseothricin resistance gene. The double deletion mutant (2KO) encodes for an additional geneticin resistance gene. As a result, in the presence of nourseothricin, each mutant ratio increased, with the disappearance of the wt virus after 5 passages (Fig 1C). These data allowed competition assays to be validated as an effective tool to assess the fitness of recombinant viruses. In the absence of selection, the relative abundance of the mutants compared to wt remained around 0.5 over 5 passages, supporting the absence of a fitness cost, even for the double deletion mutant (Fig 1C).

### Composition of the genomic fiber of single mutants

To determine the composition of the genomic fiber of each single mutant we extracted and purified their genomic fiber. MS-based proteomics confirmed that the most abundant protein was the remaining GMC-oxidoreductase. Negative staining transmission electron microscopy (NS-TEM) highlighted surprising differences between the two structures (Fig 2), despite the fact that the two proteins (qu_946 and qu_143) share 69% sequence identity (81% similarity). Specifically, the genomic fiber extracted from the KO_qu143 mutant and made of qu_946 (the most abundant in the wild type genomic fiber) was mostly in the unwound state (Fig 2B), while the one made of qu_143 (KO_qu946), resulted in very long and stable helices that did not unwind (Fig 2C). Thus, the use of both GMC-oxidoreductases in the wt genomic fiber could contribute to a fine tuning of its biophysical properties, with an intermediate state between the wt and each deletion mutant (Fig 2A). In the case of the double mutant, the protocols for capsid opening to extract the genomic fiber of wt or single mutants did not work properly. An optimized protocol allowed the extraction of a possible thinner genomic fiber, but in poor yield, preventing its purification and compositional characterization (Fig 2D).

**Figure 2:**
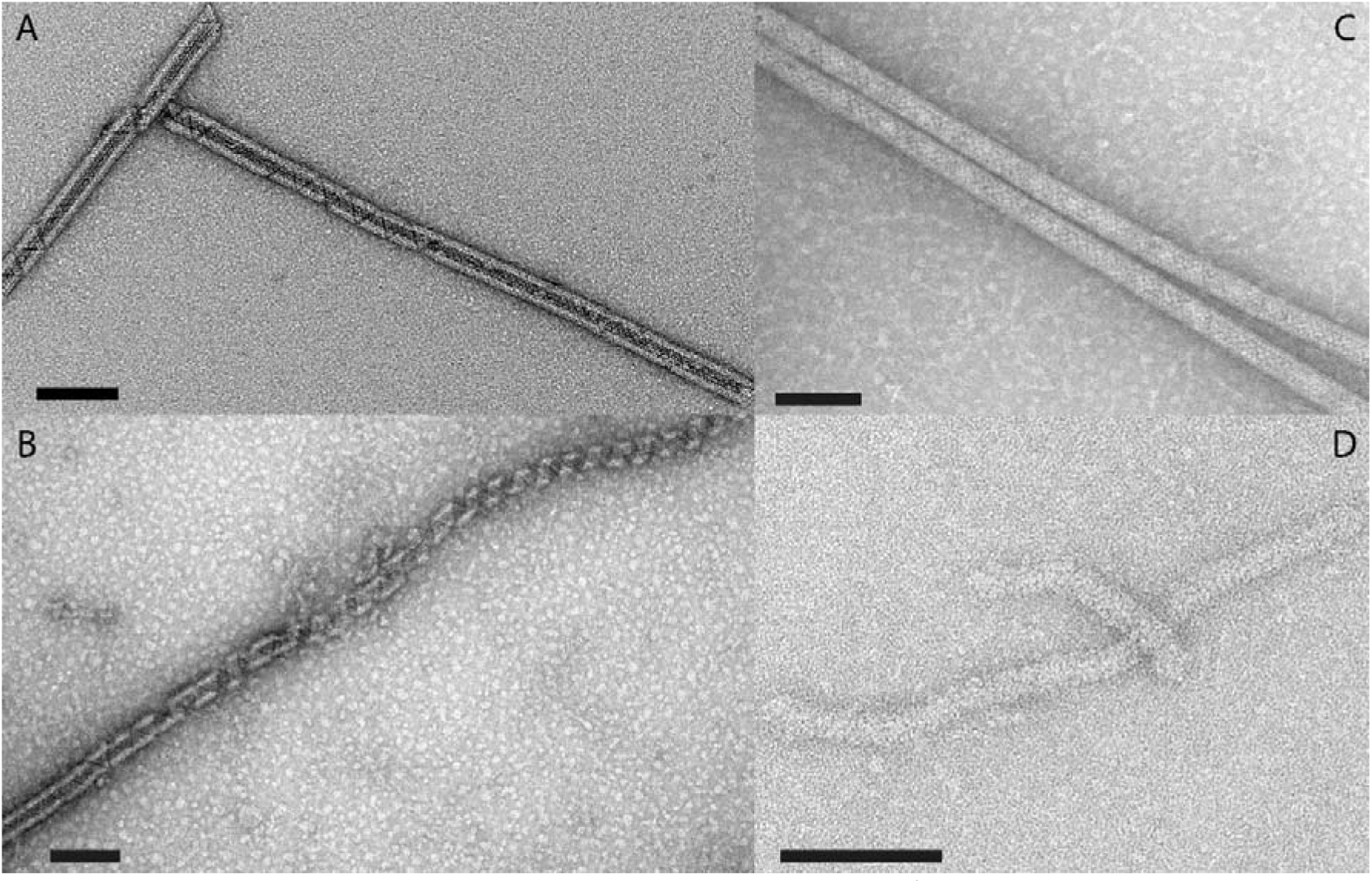
**Micrograph of negative stained genomic fiber** of A] Wild-type mimivirus, B] KO_qu143 mutant, C] KO_qu946 mutant, and D] 2KO double mutant. Scale bars 100 nm.

### Cryo-EM single particle analysis of the qu_143 genomic fiber

To determine the contribution of each GMC-oxidoreductase to the wt genomic fiber we performed cryo-EM single-particle analysis on the most stable genomic fiber composed by qu_143 (mutant KO_qu946). As for the wt, the 2D-classification revealed an heterogeneity of the sample, and the 2D classes were sorted by applying our already described clustering protocol (1). The two main clusters corresponding in width to the compact (Cl1a) and (Cl2) structures of the wt genomic fiber (Fig 3) were respectively named Cl1 and Cl2.

**Figure 3:**
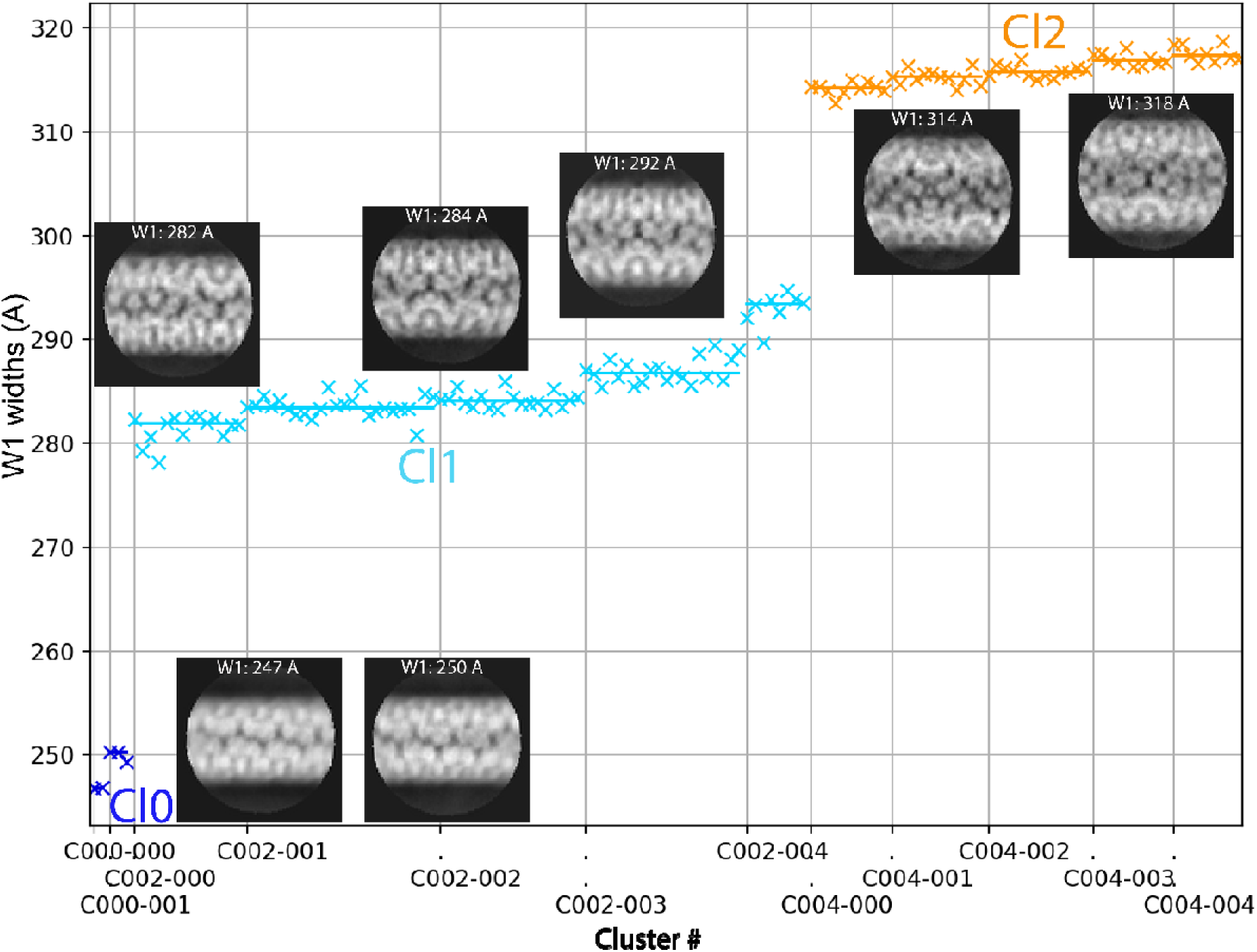
Clustering of the 2D classes obtained with qu_143 genomic fiber: Automatic sorting of the 2D classes using the fiber width W1 and pairwise correlations of the 2D classes resulted in 2 main clusters (5-start Cl1 in cyan; 6-start Cl2 in orange) and a smaller cluster (Cl0 in dark blue). Each cross corresponds to a 2D class and its associated W1. Representative 2D classes are displayed with their respective W1.

For each main cluster, we confirmed that the helical symmetry parameters were the same as the wt genomic fiber and proceeded to structure determination and refinement (Fig 4). For the less populated smaller cluster (Cl0, ∼25 nm), absent from the wt genomic fiber 2D-classes, we failed to identify its helical parameters due to the lower number of segments and the resulting lower resolution. After 3D-refinement, we obtained a 4.3 Å resolution helical ∼29 nm diameter structure (FSC threshold 0.5, masked) for Cl1. This structure corresponds to the same 5-start left-handed helix as the wt (Cl1a), made of a ∼8 nm-thick proteinaceous external shell with 5 dsDNA strands lining the interior of the shell and a ∼9 nm wide central channel (Fig 4, Fig. S2). The 4.2 Å resolution Cl2 map obtained after 3D refinement (Fig 4, Fig. S 2) corresponds to the same ∼32 nm diameter 6-start left-handed helix as the wt, with 6 dsDNA strands lining the external shell and a ∼12 nm wide inner channel (Fig 4).

**Figure 4:**
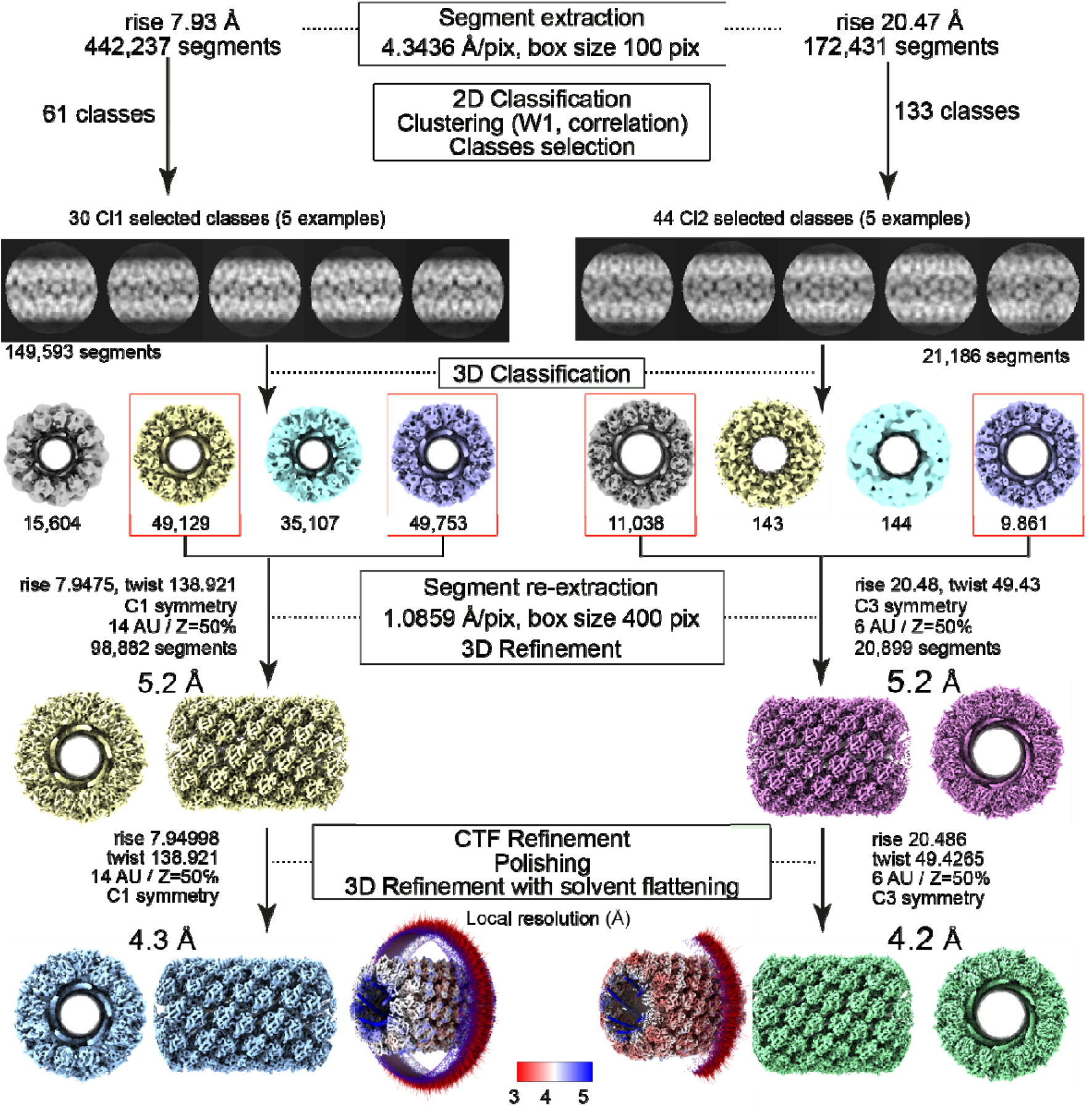
Workflow of the 5- and 6-start helices reconstruction processes. Segment extraction was performed with a box size of 400 pixels (pix) binned (box size 100 pix, 4.3436 Å/pix). The distance between consecutive boxes was equal to the axial rise calculated by indexation of the power spectrum. After clustering, 2D classes were selected for Cl1 and Cl2 and 3D classification was carried out using the selected segments, helical symmetry parameters from the power spectrum indexation, and a 300 Å or 330 Å featureless cylinder as 3D reference for Cl1 and Cl2, respectively. 3D-refinement of the 2 boxed 3D classes was achieved using one low pass filtered 3D class as reference on the unbinned segments. A first 3D-Refinement was performed with solvent flattening followed by CTF refinement and polishing of the selected segments. A last 3D refinement was achieved with solvent flattening. The EM maps colored by local resolution from 5 Å (blue) to 3 Å (red) with Euler angle distribution of the particles used for the 3D reconstruction are presented.

Since the helical parameters between the wt Cl1a and the mutant Cl1 are the same, we used the qu_143 dimeric structure refined in the Cl1a focus refined map for refinement into the mutant maps (Material and Methods section and Table S3). The density that can be attributed to the FAD cofactor was present in both maps (Fig. S3) and the models of Cl1 and Cl2 dimers are superimposable with a core RMSD of 0.467 Å based on Cα atoms (Table S4). As the qu_143 helices are more stable than the wt (composed of both GMC-oxidoreductases), the relaxed Cl3 cluster was never observed with this mutant.

### Model explaining the co-existence of 5- and 6-start helices

As for the wt, the genomic fiber of KO_qu946 is composed of a mixture of 5 and 6 strands of DNA, despite the presence of a single GMC-oxidoreductase into the shell. We estimated the ratio of 5-start and 6-start from the clustering (Fig 3) and can now propose a model that reconciles the co-occurrence of the two structures. In this model, the whole genome would be folded into 6 parallel strands, 5 longer than the 6^th^ one. The helix would then be formed initially as a 6-start helix until the sixth strand ends and, from that point, becomes a 5-start helix (Fig 5). According to this model, assuming the length of the genomic fiber is limited by the size of mimivirus nucleoid compartment, we can estimate that the maximum genome length would be ∼1.4 Mb for a full 6-start helix with an 8 nm thick protein shell. We hypothesize that the last cluster (Cl0, Fig 3) could correspond to a 4-start with an additional shorter DNA strand.

**Figure 5:**
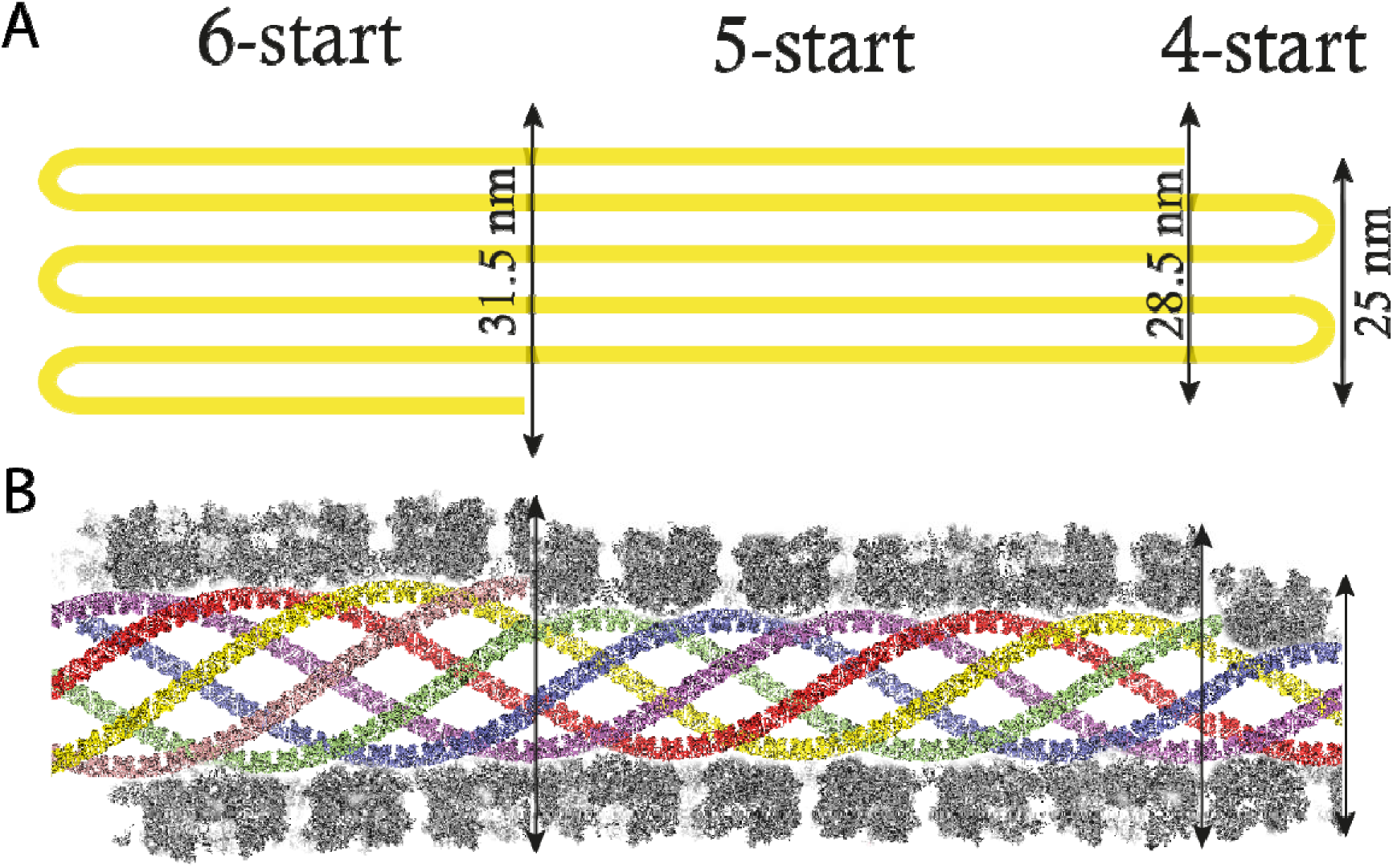
Model explaining the transition from a 6- to a 5-start helix. A] Flat models of the transition from a 6- to a 5-starts involving a decrease of the helix diameter by ∼3.5 nm. The small cluster could thus correspond to a 25 nm diameter 4-start helix. B] Cartoon model of the different helices. A longitudinal section of the GMC-oxidoreductase shell is represented around the central channel and each helix was positioned to produce a certain continuity of the DNA strands.

### The N-terminal cys-pro-rich domain is an addressing domain to the virion surface

The sequence of the cys-pro-rich N-ter domain of the GMC-oxidoreductases is not covered by proteomic analysis of the purified genomic fiber but is covered by peptides in the purified fibrils forming the external layer at the surface of the capsids (1). To assess whether this cys-pro-rich N-ter domain could be a structural signature used to address proteins to the layer of fibrils, we replaced the second GMC-oxidoreductase (qu_143) in the genome of the mutant KO_qu946 by the sequence of the GFP in fusion with the sequence of the qu_143 N-terminal domain (Nqu143-GFP). We then analyzed the resulting virions by MS-based proteomics (Table 1 and Table S2) and fluorescence microscopy. Purified Nqu143-GFP virions showed a strong fluorescence supporting the incorporation of the chimeric protein into the viral particles (Fig 6). Moreover, after defibrillation of the virions the GFP fluorescence was lost. In addition, Nqu143-GFP was identified in purified virions (ranked 97^th^ in terms of relative abundance, Table S2) and was found enriched 12-times in the fraction containing the external fibrils (ranked 20^th^, Table 1 and Table S2) with peptides covering the N-terminal domain (Fig. S5). Taken together, these data indicate that the N-terminal cys-pro-rich domain of the GMC oxidoreductase is sufficient to direct the proteins to the layer of fibrils at the surface of mimivirus particles.

**Figure 6:**
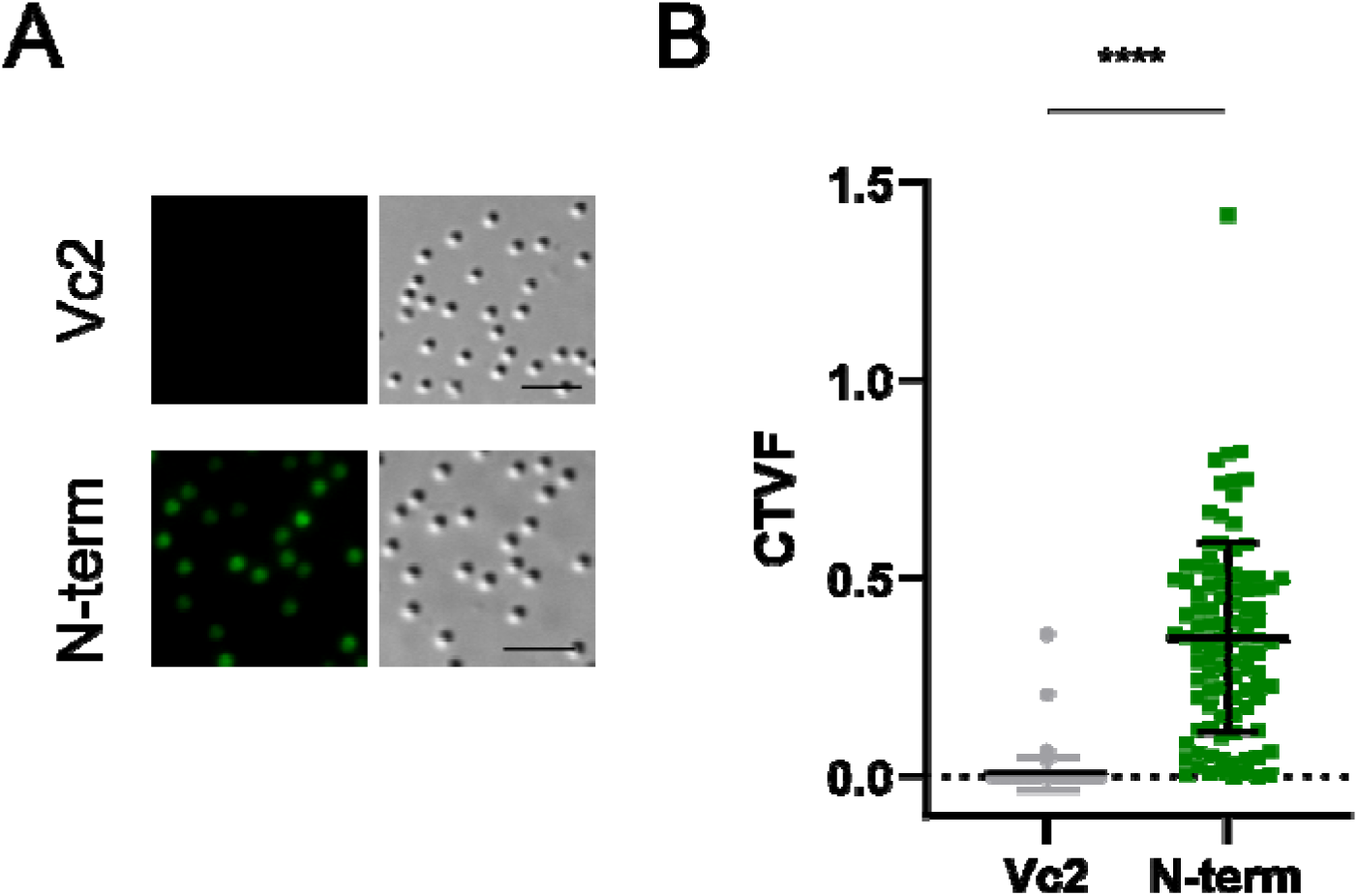
**Quantification of fluorescent virion** in infected cells overexpressing GFP (Vc2) and in cells infected by the mutant Nqu143-GFP. Scale bar 5 µm. (CTVF: corrected total virion fluorescence).

**Table 1:** Rank of the most abundant proteins in the purified fibrils of members of clade A, B and C. NA: Not applicable, the corresponding gene is absent in the corresponding genome. Sequences of the relevant orthologues of mimivirus reunion in other viruses are provided in Fig. S6.

| Rank | Mimivirus<br>reunion | KO_946 | KO_143 | 2KO | 2KO-GFP | M4 | Megavirus<br>chilensis | Megavirus<br>vitis | Moumouvirus<br>ausraliensis | Moumouvirus<br>maliensis |
| --- | --- | --- | --- | --- | --- | --- | --- | --- | --- | --- |
| qu_143 | 1 | 1 | NA | NA | NA | NA | 73 | 46 | 45 | 130 |
| qu_757 | 2 | 3 | 2 | 2 | 11 | 7 | 44 | 24 | 7 | 19 |
| qu_734 | 3 | 7 | 6 | 1 | 8 | 1 | 7 | 14 | 14 | 16 |
| qu_482 | 4 | 42 | 45 | 4 | 1 | 54 | 42 | 90 | 9 | 42 |
| qu_880 | 5 | 13 | 13 | 11 | 22 | NA | 6 | 18 | 18 | 24 |
| qu_384 | 6 | 4 | 8 | 3 | 2 | 2 | 9 | 4 | 13 | 18 |
| qu_773 | 14 | 8 | 4 | 10 | 9 | 3 | 8 | 27 | 5 | 8 |
| qu_657 | 15 | 5 | 3 | 27 | 6 | 27 | 1 | 7 | 4 | 2 |
| qu_600 | 16 | 6 | 5 | 7 | 3 | 4 | 10 | 1 | 3 | 4 |
| qu_738 | 17 | 15 | 23 | 12 | 5 | 5 | 3 | 2 | 1 | 3 |
| qu_752 | 35 | 17 | 22 | 55 | 4 | 21 | NA | NA | NA | NA |
| qu_582 | 39 | 11 | 7 | 43 | 45 | 28 | 26 | 55 | 33 | 39 |
| qu_465 | 10 | 2 | 1 | 6 | 18 | 6 | 17 | 42 | 32 | 29 |
| mm_751 | NA | NA | NA | NA | NA | NA | 2 | 3 | 2 | 1 |
| Nqu143-GFP | NA | NA | NA | NA | 20 | NA | NA | NA | NA | NA |

### Composition of the purified external fibrils

#### Protein composition of the layer of fibrils in additional members of the family Mimiviridae

The members of the *Megavirinae* subfamily (20) currently encompasses five clades, the mimiviruses (A clade), moumouviruses (B clade), megaviruses (C clade), tupanviruses (D clade) (8) and cotonvirus japonicus (E clade) (21). As of today, all members of the *Megavirinae* are characterized by a layer of fibrils that differs in thickness and lengths among the clades and, excepted for clade A (1, 18), their protein composition was unknown. While the cys-pro-rich N-ter domain of the GMC-oxidoreductases is conserved in all members of the A clade, it is absent in the orthologous proteins in members of the B and C clades. Moreover, homologs of the two GMC-oxidoreductases are pseudogenized in tupanviruses, suggesting that different proteins should compose their fibrils (Fig. S6). We thus conducted a systematic proteomic analysis of the purified fibrils of different members of the 3 clades, in addition to the mutants. For the double mutant (2KO) we identified a group of proteins as the most abundant in the fibrils fraction (qu_734, qu_757, qu_384 and qu_482, Table 1 and Table S2). These proteins were also highly ranked in the Nqu143-GFP double mutant fibrils and the orthologous of qu_734 was identified as the most abundant protein in the laboratory-evolved M4 mutant fibrils (696-L688, Table 1 and Table S2). This mutant does not encode a glycosylation machinery and is not glycosylated (4, 18), thus its capsid lacks the large reticulated layer of fibrils decorating mimivirus capsid (Fig. S8F). Interestingly the qu_734 protein also possesses a cys-pro-rich N-ter domain which is conserved in all clades (Fig. S6). It was not possible to determine if one of these proteins was also the building block composing the 2KO genomic fiber, given the difficulty to open the 2KO capsids. Thus, we concluded that the change in protein composition of the fibrils led to capsids with different stability properties. While, as for the wt, the most abundant protein in KO_qu946 remains qu_143, the lack of qu143 in KO_qu143 does not lead to its replacement by the second GMC-oxidoreductase. Instead, it is replaced by a group of proteins, with qu_465 (predicted as a thioredoxin domain containing protein) as the most abundant, followed by qu_757, the second most abundant in the wt fibrils (Table I and Table S2).

The proteomic analysis of fibrils purified from two isolates of the B clade, moumouvirus australiensis and maliensis, showed similar protein compositions, but with slight differences in their relative abundances (Table 1 and Table S2). The moumouvirus maliensis protein mm_751, ranked 1^st^ in its fibrils, is absent from A clade fibrils and is in the top 3 in the fibrils of members of the B and C clades. Cystein and proline amino acids are present in the N-ter domain of mm_751, but less abundant than in the GMC-oxidoreductases. For ma_195 (qu_738 in mimivirus reunion, Table 1 and Table S2), ranked 1^st^ in the fibrils of moumouvirus australiensis, the N-ter domain is not cys-pro-rich. The best ranked in the fibrils of members of the C clade are the same as for members of the B clade. The first ranked in megavirus chilensis is mg749 (qu_657 in mimivirus reunion) which presents a cys-pro-rich N-ter domain, while the first ranked in megavirus vitis, mvi_646 (qu_600 in mimivirus reunion), lacks a cys-pro-rich N-ter domain. Yet, for each virus in each clade, there is at least one protein with a cys-pro-rich N-terminal domain among the most abundant in the fibrils.

#### Glycan composition of the fibrils in mimivirus mutants

We recently established that the fibrils were coated with glycans (3, 9) and demonstrated that for mimivirus, the prototype of the A clade, they were made of two distinct large molecular weight polysaccharides (4). Since the cluster of 12 genes responsible for the biosynthesis of the polysaccharides is conserved in the A clade (3), we hypothesized that the fibrils of the mimivirus reunion strain would have the same composition as mimivirus. To assess whether the knockout of the most abundant protein composing the fibrils could also affect the branching of these polysaccharides and their composition, we analyzed the sugar content of the viral particles of all three mimivirus reunion mutants (KO_qu946, KO_qu143, and 2KO) together with the wt strain, as reported in (3, 4). The chemical characterization revealed for each of them the presence of sugars (Figure 7A), with rhamnose (Rha), viosamine (Vio), 2-OMe-Vio, glucose (Glc), and glucosamine (GlcN) (Fig 7A), confirming that mimivirus reunion strain and all mutants had the same glycan composition as mimivirus prototype (3). We then analyzed the fibrils of the mutants and wt by ^1^H NMR spectroscopy (22) and compared them with the mimivirus prototype, which confirmed that the different sugars were assembled the same way as in the reference mimivirus to produce the same two polysaccharides (Fig 7, Fig. S7) (4). NS-TEM images of the virions of the different mutants were obtained after methylcellulose staining to assess whether the mutations changed the layer of fibrils appearance, which would suggest that the change in protein impacted their level of glycosylation (23). All mutants’ virions showed the same cross-linked outer layer, suggesting that despite their differences in protein composition, their glycosylation was not affected (Fig. S8).

**Figure 7:**
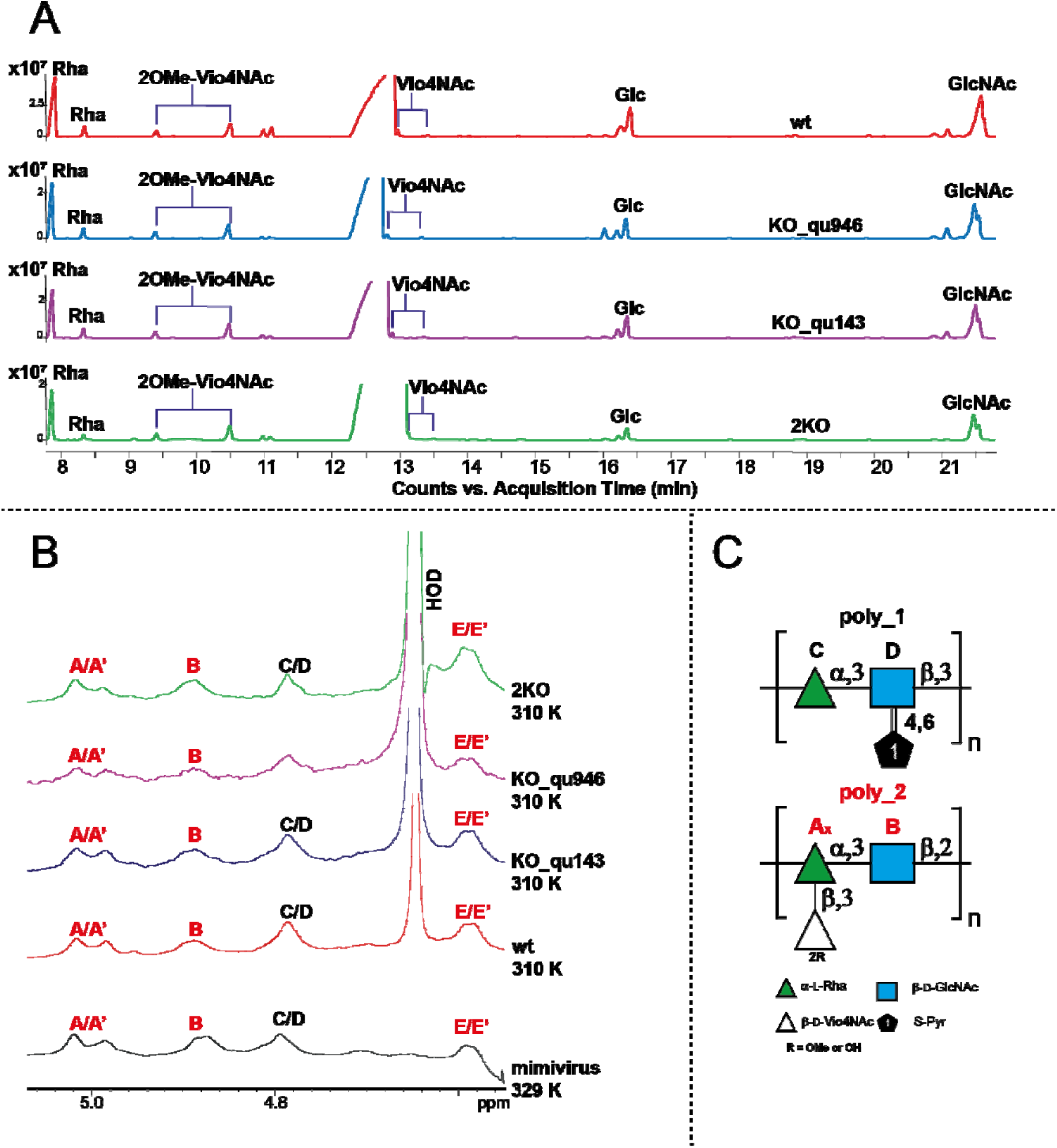
**Compositional and NMR analysis of the fibrils** of mimivirus reunion strain wt and mutants A] GC-MS chromatogram profiles of the sugars composing the fibrils of wt (a), KO_qu946 (b), KO_qu143 (c), and 2KO (d). B] Comparison of the ^1^H NMR spectra of mimivirus reunion strain wt and related mutants with that of mimivirus prototype strain. The anomeric signals related to poly_1 (**C** and **D** units, in black), and those of poly_2 are in red. C] Structures of mimivirus polysaccharides as reported in Notaro *et al*, 2021 (4).

It was hypothesized that mimivirus R135 GMC oxidoreductases (qu_143 in mimivrius reunion) was the major target for glycosylation in the surface fibrils of mimivirus (4, 18). The results presented here indicate that the mutations, including the deletion of the two GMC-oxidoreductase genes, did not affect the surface glycosylation. These data suggest that the GMC oxidoreductases are not the main target for glycosylation. Alternatively, it is also possible that the glycosylation machinery was able to use the qu_734 protein as support for the two polysaccharides. As expected, no glycans were identified in the laboratory-evolved M4 mutant layer of fibrils.

## Discussion

Genome packaging is essential for the propagation of viruses. For instance, the packaging ATPase of poxviruses (24) or the histones of marseilleviruses (25, 26) are essential for the productive infection by their virions. In addition, proteins at the surface of viral capsids usually play a central role in interacting with the host cell surface and the initial steps of infection (27). The two processes, genome encapsidation and viral entry, are thus essential. As the GMC oxidoreductase are composing both the genomic fiber and the surface fibrils (1) we could have expected them to be essential. However, these two enzymes are absent in the laboratory-evolved mimivirus M4 strain that also lost the glycosylation machinery (18). Our recent implementation of genetics tools for cytoplasmic giant viruses provided the first opportunity to directly assess the impact of the GMC-oxidoreductases deletions in mimivirus reunion strain (19). Given the homology between the two enzymes, the deletion of the most abundant one in the wt genomic fiber induced its replacement by the second one, with no apparent cost for the virus, in laboratory conditions. Yet, the single mutant genomic fibers were significantly different compared to the wt (Fig. 2), indicating a certain degree of functional specialization for each GMC-oxidoreductase. While the wt genomic fiber was highly heterogeneous, with fully relaxed helices having lost DNA (1), the KO_qu946 genomic fiber appeared more stable, with no occurrence of fully relaxed structures. In contrast, the deletion of qu_143 seemed to increase the genomic fiber instability, suggesting a stabilizing role for this second GMC-oxidoreductase (Fig. 2). The cryo-EM analysis of the KO_qu946 genomic fiber structure, as for the wt, produced the same two 5- and 6-start helices, but composed of a single GMC-oxidoreductase. This led us to propose a model to explain the co-existence of both structures. In this model, the genome is folded into six strands before assembly into a 6-start helix. The first and the last strand can be shorter than the other strands, leading to a 5-start helix with 5 DNA strands and a 4-start helix with 4 DNA strands. Finally, the deletion of both GMC-oxidoreductases with no significant loss of viral fitness demonstrated they were not essential.

The qu_143 GMC oxidoreductase is 5 times less abundant than qu_946 in the genomic fiber but it is ranked 1^st^ in the layer of fibrils and is 3 times more abundant than the next one, qu_757. It was previously predicted as the main target for glycosylation (4, 17, 18) while qu_946 is only ranked 13^th^ in the fibrils (Table S2). MS-based proteomics revealed that the cys-pro-rich N-ter domain of the GMC-oxidoreductase is present in the fibrils and cleaved in the genomic fiber (1), but the protease and precise cleavage site remains to be identified.

Interestingly, an additional mutant in which this cys-pro-rich N-ter domain was fused to GFP, produced virions in which the GFP was identified at the surface of the virions and is enriched in the surface layer of fibrils. Further studies will be needed to elucidate the mechanism behind the recognition of this domain and its addressing to the fibrils. In addition, the virions of all mutants were decorated by long, reticulated fibrils as evidenced by TEM (Fig. S8).

Except for the KO_qu946 mutant, MS-based proteomic analysis of the fibrils revealed they were composed by other proteins some of which also presenting a cys-pro-rich N-ter domain (Fig. S6). Finally, the glycan analysis of the GMC-oxidoreductases single and double mutants confirmed that their fibrils were still glycosylated by the same two polysaccharides as the wt virions.

In a previous study, it was reported that the glycosylation machinery was clade-specific and produced different glycans (3). In the present study, the analysis of the purified fibrils of members of the B and C clades revealed they were also composed by different proteins inter- and even intra-clade and that B and C clades virions’ fibrils present a protein composition closer to each other than that of A clade.

The glycosylated layer of fibrils bear glycans echoing bacterial ones, recognized by the amoeba which feeds on bacteria (2, 3). They thus appear as key for productive infection. In a given environment, the variable composition of the fibrils could be used to favor the engulfment of a given virion by a given amoeba, compared to others virions and even bacteria. The protein content of the layer of fibrils appears to have been optimized in a given clade, but their complex composition suggests they can be made from a large set of diverse proteins. Having a flexible toolbox for building the external layer of fibrils would reflect the need to constantly adjust the capsid composition to outcompete other parasites and secure infection. We can thus hypothesize that in the population of virions resulting from a single infection, this composition might also be variable, helping out the virus to ensure the productive infection of at least one of the possible hosts present in that environment by at least one virion.

The question whether it is always the same protein that makes both the layer of fibril and the genomic fiber, even when other proteins than the GMC-oxidoreductases are used, remains unanswered. Yet, the most abundant protein composing the external layer of fibrils of the 2KO and M4 corresponds to the 222 amino-acids qu_734 protein, which according to the high confidence alphaFold (28) prediction could be almost twice smaller (∼4.5 nm) than the GMC-oxidoreductase. From the previous study, the central part of the genomic fiber has to correspond to a central channel of at least 9 nm to accommodate proteins such as the RNA polymerase (1). There is 4 additional nm for the DNA lining the protein shell and the GMC-oxidoreductase is 8 nm height and makes the 16 nm helical shell, leading to a ∼29 nm helix.

The protein making the shell is thus defining the final dimension of the genomic fiber. For the 2KO, if this is the same protein that makes the external layer of fibrils and the shell of the genomic fiber, we expect a genomic fiber of 22 nm, compatible with the thinner structure observed in Fig 1D (∼20-25 nm).

We have previously proposed an increased genome redundancy as a contributing factor to the appearance of viral gigantism (29). The data presented here validate such premise and extend these predictions outside of the *Pandoraviridae*. Overall, our results reveal the resilience of mimivirus, with redundant solutions securing essential functions such as infectiousness and genome packaging. Functional redundancy, well documented in the cellular world as a way to preserve essential function such as cell division (30), and until now a hallmark of the optimized microorganisms, may thus also be at work in the viral world.

## Material and methods

### Cloning of DNA constructs

A detailed protocol for gene manipulation of giant viruses and their host is provided in (19).

#### Gene knock-out vectors

The plasmid for gene knock-out was generated by sequential cloning of the 3’ UTR of mg_18 (megavirus chiliensis), the promoter of mg_741 (megavirus chiliensis), and a Nourseothricin N-acetyl transferase (NAT) or a neomycin selection cassette (NEO). Each cloning step was performed using the Phusion Taq polymerase (ThermoFisher) and InFusion (Takara). Finally, 500bp homology arms were introduced at the 5’ and 3’ end of the cassette to induce homologous recombination with the viral DNA (19). Before transfection, plasmids were digested with EcoRI and NotI. All primers are shown in FigS1.

### Establishment of viral lines

#### Gene knock-out

Gene knockout strategy was performed as previously described for pandoravirus (29). Briefly, 1.5x105 *Acanthamoeba castellanii cells* were transfected with 6 μg of linearized plasmid using Polyfect (QIAGEN) in phosphate saline buffer (PBS). One hour after transfection, PBS was replaced with PPYG, and cells were infected with 1.5x10^7^ mimivirus reunion particles for 1 hour with sequential washes to remove extracellular virions. 24h after infection the new generation of viruses (P0) was collected and used to infect new cells. An aliquot of P0 viruses was utilized for genotyping to confirm the integration of the selection cassette. Primers used for genotyping are shown in Table S1. A new infection was allowed to proceed for 1 hour, then washed to remove extracellular virions and nourseothricin was added to the media. Viral growth was allowed to proceed for 24 hours. This procedure was repeated one more time before removing the nourseothricin selection to allow viruses to expand more rapidly. Once, the viral infection was visible, the selection procedure was repeated one more time. Viruses produced after this new round of selection were used for genotyping and cloning (19). Double knockout of the GMC oxidoreductases was obtaining by using a clonal population of qu_143 knockout viruses as parental strain. The locus of qu_946 was replaced by a neomycin resistance cassette. The transfection and selection of recombinant viruses’ procedure was performed identical to the process to generate single knockout but replacing nourseothricin by geneticin.

#### Cloning and genotyping

150,000 *A. castellanii* cells were seeded on 6 well plates with 2 mL of PPYG. After adhesion, viruses were added to the well at a multiplicity of infection (MOI) of 1. One-hour post-infection, the well was washed 5 times with 1mL of PPYG, and cells were recovered by well scraping. Amoebas were then diluted until obtaining a suspension of 1 amoeba/μL. 1μL of such suspension was added in each well of a 96-well plate containing 1000 uninfected *A. castellanii* cells and 200 μL of PPYG. Wells were later monitored for cell death and 100 μL collected for genotyping (19). Genotyping was performed using Terra PCR Direct Polymerase Mix (Takara) following the manufacturer’s specifications. Primers used for genotyping are shown in Table S1.

### Competition assay and quantitative PCR analysis

Equal infectious particles of wild-type and recombinant mimivirus reunion were mixed and used to infect *A. castellanii* at an approximate multiplicity of infection of 0.8. Viruses were allowed to grow overnight in the presence or absence of nourseothricin. Subsequent viral progenies were used to infect new *A. castellanii* cells in reiterative passages. A fraction of each passage was collected for genomic DNA extraction.

Viral genomes were purified using Wizard genomic DNA purification kit (PROMEGA). To determine the amplification kinetic, the fluorescence of the EvaGreen dye incorporated into the PCR product was measured at the end of each cycle using SoFast EvaGreen Supermix 2× kit (Bio-Rad, France). A standard curve using the gDNA of purified viruses was performed in parallel with each experiment. For each point, a technical triplicate was performed. Quantitative real-time PCR (qRT-PCR) analyses were performed on a CFX96 Real-Time System (Bio-Rad).

### Genome sequencing and assembly of mutants’ genomes

Genomic DNA was extracted from 10^10^ virus particles using the PureLink TM Genomic DNA mini kit (Invitrogen) according to the manufacturer’s protocol. Clones of individual mutants and wt were sequenced on an illumina platform (Novogen). For mimivirus wt, we obtained 4,819,885 150nt paired-end reads. For KO_qu946, KO_qu946 and 2KO, 4,662,744, 4,572,873 and 5,068,030 150nt paired-end reads, respectively. Genomes were assembled using spades v3.13, with the option “careful” and we obtained 5 contigs for the wt, 4 for KO_qu946, and 7 for KO_qu143 and 2KO. The wt sequence was consistent with the original mimivirus reunion genome sequence (GI MW004169) and for the clone KO_qu946 the gene was interrupted from position 1,151,358 to 1,152,985 relative to wt genome (position 302 to 1741 in the corresponding qu_946 gene) and for the clone KO_qu143 the gene was interrupted from position 165,934 to 167,572 relative to wt genome (position 306 to 1741 in the corresponding qu_143 gene). The region is replaced in both cases by the nourseothricin cassette sequence (1637 nt). For the 2KO, the gene qu_143 is interrupted from positions 306 to 1741 (165,934 to 167,572 on the wt genome) and replaced by the nourseothricin cassette sequence (1637 nt), and gene qu_946 is interrupted from positions 302 to 1206 (1,151,551 to 1,153,410 on the genome) and replaced by the geneticin cassette sequence (1859 nt). To confirm the deletion of each mutant, the reads were mapped on the wt genome resulting in homogeneous coverage along the genome, except for the qu_143 and qu_946 central positions which are covered. In addition, we used the central part of the GMC-oxidoreductase genes (deleted in mutants) as blast queries against the reads of each mutant genome, which also confirmed the absence of the central region.

### Extraction and purification of the qu_946 and qu_143 mutants’ genomic fiber

The genomic fiber of the mimivirus reunion single mutants of qu_946 (KO_qu946) and qu_143 (KO_qu143), were extracted as described in Villalta et al, 2022 for the wt virus (1). The genomic fiber was extracted from 12 mL of purified single deletion mutant virions at 2 x 10^10^ particles/mL, split into 12 x 1 mL samples processed in parallel. Trypsin (Sigma T8003) in 40 mM Tris-HCl pH 7.5 buffer was added at a final concentration of 50 µg/mL and the virus-enzyme mix was incubated for 2h at 30°C in a heating dry block (Grant Bio PCH-1). DTT was then added at a final concentration of 10 mM and incubated at 30°C for 16h. Finally, 0.001% Triton X-100 was added to the mix and incubated for 4h at 30°C. Each tube was centrifuged at 4,000 x g for 5 min to pellet the opened capsids. The supernatant was recovered and concentrated by centrifugation at 15,000 x g for 4h at 4°C. Most of the supernatant was discarded leaving 12x∼200 µL of concentrated broken pieces of genomic fiber that were pooled and layered on top of ultracentrifuge tubes of 4 mL (polypropylene centrifuge tubes, Beckman Coulter) containing a discontinuous cesium chloride gradient (1.4, 1.3, 1.2 g/cm^3^ in 40 mM Tris-HCl pH 7.5 buffer). The gradients were centrifuged at 200,000 x g for 16h at 4 °C. Since no visible band was observed, successive 0.5 mL fractions were recovered from the bottom of the tube. Each fraction was dialyzed using 20 kDa Slide-A-Lyzers (ThermoFisher) against 40 mM Tris-HCl pH 7.5 to remove the CsCl. These fractions were further concentrated by centrifugation at 15,000 x g, at 4°C for 4h, and most of the supernatant was removed, leaving ∼100 µL of sample at the bottom of each tube. At each step of the extraction procedure, the sample was imaged by negative staining transmission electron microscopy (NS-TEM) to assess the integrity of the genomic fiber. Each fraction of the gradient was finally controlled by NS-TEM.

### Negative staining TEM

300 mesh ultra-thin carbon-coated copper grids (Electron Microscopy Sciences, EMS) were prepared for negative staining by adsorbing 4-7 µL of the sample for 3 min., followed by blotting excess liquid and staining for 2 min in 2% uranyl acetate to image the genomic fiber. For fibrils and mutant virions, staining was performed with a drop of 1% uranyl followed by blotting after 10-15 s, and a drop of uranyl acetate coupled with methylcellulose (2% and 0.2%, respectively) was added twice and left for 5-10 s before blotting.

The grids were imaged either on an FEI Tecnai G2 microscope operated at 200 keV and equipped with an Olympus Veleta 2k camera (IBDM microscopy platform, Marseille, France); an FEI Tecnai G2 microscope operated at 200 keV and equipped with a Gatan OneView camera (IMM, microscopy platform, France).

### Single-particle analysis by cryo-EM

#### Sample preparation

For the KO_qu946 mutant, 3 µL of the purified sample were applied to glow-discharged Quantifoil R 2/1 Cu grids, blotted for 2 s using a Vitrobot Mk IV (Thermo Scientific) and applying the following parameters: 4°C, 100% humidity, blotting force 0, and plunge frozen in liquid ethane/propane cooled to liquid nitrogen temperature.

#### Data acquisition

Grids were imaged using a Titan Krios (Thermo Scientific) microscope operated at 300 keV and equipped with a K3 direct electron detector and a GIF BioQuantum energy filter (Gatan). 2,224 movie frames were collected using the EPU software (Thermo Scientific) at a nominal magnification of 81,000x with a pixel size of 1.0859 Å and a defocus range of -0.6 to -2.8 μm. Micrographs were acquired using EPU (Thermo Scientific) with 2.3 s exposure time, fractionated into 40 frames, and 18.25 e^-^/pixel/ s (total fluence of 35.597 e^-^/Å²).

#### 2D classification and clustering of 2D classes

All movie frames were aligned using MotionCor2 (31) and used for contrast transfer function (CTF) estimation with CTFFIND-4.1 (32). Helical segments of the purified genomic fiber manually picked with Relion 3.1.0, were extracted with 400 pixels box sizes (decimated to 100 pixels) using a rise of 7.93 Å and a tube diameter of 300 Å. Particles were subjected to reference-free 2D classification in Relion 3.1.0 (33, 34). We then performed additional cluster analysis of the initial 2D classes provided by Relion to aim for more homogeneous clusters (1), eventually corresponding to different states (Fig. 3 & 4).

#### Identification of candidate helical parameters

Fourier transform analysis methods have been used to confirm the helical parameters were the same as in wt (1, 35–37) for the Cl1 (Cl1a in wt) and Cl2.

#### Cryo-EM data processing and 3D reconstruction

After helical parameters determination, segments were extracted with a box size of 400 pixels (decimated to 100 pixels) using the proper rises for the Cl1 (7.93 Å, cylinder 300 Å, 442,237 segments) and the Cl2 (20.47 Å, cylinder 330 Å, 172,431 segments). A dedicated 2D classification protocol was performed independently on each extraction. For the Cl2, one round of 50 expectation-maximization (E-M) iterations was sufficient to produce 133 homogeneous 2D classes submitted to cluster analysis and 44 2D classes were selected (21,186 segments). For the Cl1 extraction, three iterative 2D classification/selection rounds were performed (25, 50 and 100 E-M iterations) producing 61 classes from which 30 (149,593 segments) were finally selected based on the cluster analysis.

Values of the helical parameters, rise and twist (Cl1: 7.9475 Å, 138.921°; Cl2: 20.48 Å, 49.43°), were then used for Relion 3D classification (33, 34), with a +/-0.5 units freedom search range, using a featureless cylinder as initial reference (diameter of 300 Å and C1 symmetry for Cl1 and 330 Å and C3 symmetry for Cl2). In real space, the helical symmetry was searched using 50% of the central part of the box for both structures. The number of asymmetrical units in each segment box was set to 1 for Cl1 and 6 for Cl2. The entire helical reconstruction and protein shell dimensions were obtained using an in-house developed program.

The superimposable 3D classes (same helical parameters, same helix orientation) were then selected, reducing the data set to 98,882 segments for the 5-start helix (Cl1) and 20,899 segments for the 6-start helix (Cl2). After re-extraction of the selected segments without scaling, a first unmasked 3D refinement was performed with a rescaled 3D classification output low pass filtered to 15 Å as reference, followed by a 3D refinement step using solvent flattened, FSCs and CTF refinement using the standard procedure described in Relion. To further improve the resolution of the maps, Bayesian polishing was applied using 10,000 segments for training and default values for polishing. A last round of 3D refinement with solvent flattening was applied to the previous refined map using the polished particles. At that point, the maps were resolved enough (Cl1: 4.3 Å, Cl2: 4.2; FSC threshold 0.5) to identify secondary structure elements (with visible cylinders corresponding to the helices) and were used to compute local resolution.

#### Structures refinement

The best resolution Cl2 map was used to fit the qu_143 dimeric structure (PDB 7YX3) using UCSF ChimeraX 1.5 (38). Each monomer was then rigid-body fitted independently into the map. The entire protein was then inspected within coot 0.9.7 (39) to fix local inconsistencies and was further refined against the map using the real-space refinement program in PHENIX 1.20.1 (40). The protein was submitted to 5 cycles of rigid body refinement (with each chain defined) followed by twice 10 cycles of refinement with the default PHENIX options. The resulting structure was manually corrected within coot. The resulting protein model was submitted to the same steps refinement in PHENIX. This final model was then fitted into the Cl1 map, inspected with coot and refined using 5 cycles of rigid body refinement and simulated annealing followed by twice 10 cycles of refinement with the default PHENIX options. Validations were performed with PHENIX using the comprehensive validation program (Table S3). RMSD between different structures (monomers and dimers, Table S4) were computed using the align procedure in Pymol suite (Schrödinger, L., & DeLano, W. (2020). *PyMOL*).

### Extraction and purification of the mutants’ fibrils

To analyze the glycan composition and polysaccharides structures of the 3 mutants and wt mimivirus reunion strain we applied an already described protocol (4). Briefly, 4 x 10^11^ viral particles were centrifuged at 14,000g for 10 min, the supernatant was discarded and the pellet was re-suspended in 10 ml of 50 mM DTT. Fibril extraction was performed at 100°C under stirring. After 2 h, the tube was centrifuged at 14,000 g for 15 min, at 4°C, and the fibrils recovered with the supernatant. The fibrils were then dried and purified on Biogel P10, followed by subsequent NMR analysis of each mutant and wt.

We also developed a softer defibrillation protocol to recover the fibrils without contaminating them with proteins released by damaged virions in order to analyze the fibrils protein composition by MS-based proteomics. Purified virions (1.5 x 10^10^) were incubated in Eppendorf tubes in 500 µL 40 mM Tris-HCl pH 7.5 buffer, 500 mM DTT for 2h at 30°C. Tubes were then centrifuged at 14,000 g for 10 min. The supernatants containing the fibrils were recovered and concentrated on vivaspin® 3 KDa (Sartorius, VS04T91) at 3,000 g. The pellet was washed twice with 40 mM Tris-HCl pH 7.5 buffer and centrifuged at 14,000 g for 10 min and finally resuspended in the same buffer. Intact virions, pellets and fibrils were imaged by NS-TEM to assess the integrity of the defibrillated virions in the pellet and the presence of fibrils in the supernatant. For the Nqu143-GFP mutants, defibrillated virions were also observed by fluorescence microscopy which confirmed the absence of fluorescence due to the removal of the GFP together with the layer of fibrils.

### Sugar composition of viral particles of Mimivirus reunion wt and mutants

Monosaccharide composition analyses as acetylated methyl glycoside (AMG) were performed on the intact viral particles (1,25 x 10^10^, ∼250 μl) of mimivirus reunion wt and mutants, following the procedure reported by De Castro *et al* 2010 (41). The obtained AMG were analyzed via gas chromatography-mass spectrometry (GC-MS) on an Agilent instrument (GC instrument Agilent 6850 coupled to MS Agilent 5973) equipped with a SPB-5 capillary column (Supelco, 30 m × 0.25 i.d., flow rate, 0.8 mL min–1) and He as the carrier gas. The identification of the monosaccharides derivatized as AMG, was obtained by studying the fragmentation pattern corresponding to each peak of the chromatogram and by comparison with suitable standards.

### Purification and ^1^H NMR analysis of the fibrils

The fibrils of mimivirus reunion wt and mutants, extracted as reported above, were purified to remove the DTT used for the extraction procedure.

Briefly, the glycoproteins (protein/s carrying the polysaccharides) were precipitated with cold acetone at 80%, at 4 °C, for 16 hours, twice. The supernatant containing DTT and salts was discarded, while the precipitate was dissolved in water and freeze-dried. Then, the precipitate was purified by size exclusion chromatography (Biogel P10, flow: 12 ml / h) to completely remove the DTT. The eluted fractions were checked by ^1^H NMR, revealing that the glycan-containing material was eluted at one-third of the column volume (full spectra are shown in Fig 7B).

The ^1^H NMR measurements were carried out on 600 MHz Bruker instrument, equipped with a CryoProbe™ at 310 K. The intensity of the solvent signal was reduced by measuring a mono-dimensional DOSY spectrum, setting δ and Δ to 2.4 ms and 100 ms, respectively, and the variable gradient to 50% of its maximum power. Spectra were processed and analyzed using Bruker TopSpin 4.0.9 program.

In the ^1^H NMR spectra, the anomeric region (22) (5.5-4.4 ppm) perfectly overlapped with that of the previously studied mimivirus strain (4) (Fig 7B, Fig 7-figure supplement 7), confirming that the different sugars were assembled the same way as in the reference mimivirus to produce the same two polysaccharides (Fig 7C). The signal at 4.80 ppm is the result of the overlap of two anomeric protons, the one of a 3-α-Rha (labeled **C**) and of a 3-β-GlcNAc (**D**) modified with a pyruvic acid at the hydroxyl functions 4 and 6. These two residues are the building blocks of the repeating unit of polysaccharide 1 (**poly_1**, **Fig 7C**) (4). The other anomeric signals, labeled with the capital letters **A, A’** (2,3-α-L-Rha), **B** (3-β-D-GlcNAc), **E** (2OMe-β-D-VioNAc) and **E’**(β-D-Vio4NAc), are part of the polysaccharide 2 (**poly_2**) that presents a backbone with a disaccharide repeating unit of 2)-α-L-Rha-(1→3)-β-D-GlcNAc-(1→, with the rhamnose residue further substituted with a viosamine methylated at position 2 (**E**) or not methylated (**E’unit**), thus taking the labels **A** and **A’**, respectively (Fig 7C) (4).

### Mass spectrometry-based proteomic analyses

Proteins were solubilized with Laemmli buffer (4 volumes of sample with 1 volume of Laemmli 5X - 125 mM Tris-HCl pH 6.8, 10% SDS, 20% glycerol, 25% β-mercaptoethanol and traces of bromophenol blue) and heated for 10 min at 95 °C. The extracted proteins were stacked in the top of an SDS-PAGE gel (4-12% NuPAGE, Life Technologies), stained with Coomassie blue R-250 (Bio-Rad) before in-gel digestion using modified trypsin (Promega, sequencing grade) as previously described (42). The resulting peptides were analyzed by online nanoliquid chromatography coupled to tandem MS (UltiMate 3000 RSLCnano and Q-Exactive Plus or Q-Exactive HF, Thermo Scientific). Peptides were sampled on a 300 µm x 5 mm PepMap C18 precolumn and separated on a 75 µm x 250 mm C18 column (Reprosil-Pur 120 C18-AQ, 1.9 μm, Dr. Maisch, except for KO_qu143, KO_qu946 and Nqu143-GFP mutant samples separated on Aurora, 1.7 µm, IonOpticks) using a 140-min gradient (except for fibrils from Nqu143-GFP mutant for which a 60-min gradient was used). MS and MS/MS data were acquired using Xcalibur 4.0 (Thermo Scientific). Peptides and proteins were identified using Mascot (version 2.8.0, Matrix Science) through concomitant searches against homemade *A. castellanii* protein sequence database, homemade virus-specific protein sequence databases, and a homemade database containing the sequences of classical contaminant proteins found in proteomic analyses (human keratins, trypsin…). Trypsin/P was chosen as the enzyme and two missed cleavages were allowed. Precursor and fragment mass error tolerances were set at respectively at 10 and 20 ppm. Peptide modifications allowed during the search were: Carbamidomethyl (C, fixed), Acetyl (Protein N-term, variable) and Oxidation (M, variable). The Proline software version 2.2.0 (43) was used for the compilation, grouping, and filtering of the results: conservation of rank 1 peptides, peptide length ≥ 6 amino acids, peptide-spectrum-match identification false discovery rate < 1% (44), and minimum of 1 specific peptide per identified protein group. Proline was then used to perform a MS1-based quantification of the identified protein groups. Intensity-based absolute quantification (iBAQ) (45) values were calculated for each protein group from the MS intensities of razor and specific peptides (Table S2). The relative abundance of individual proteins in virions and fibrils was calculated as the ratio of the individual protein iBAQ values to the sum of the iBAQ values of all proteins in each sample. The relative enrichment of individual proteins between virions and fibrils was calculated as the ratio of their relative abundances in each fraction (1).

## Acknowledgments

We thank Jean-Michel Claverie for his comments on the manuscript and discussions all along the project and Elsa Garcin for editing the revised manuscript. The data were collected at the Cryo-EM Swedish National Facility funded by the Knut and Alice Wallenberg, Family Erling Persson and Kempe Foundations, SciLifeLab, Stockholm University, and Umeå University under the expert assistance of Dr. Matthieu Coinçon. The preliminary electron microscopy experiments were performed on the PiCSL-FBI core facility (Aïcha Aouane, IBDM, AMU-Marseille), a member of the France-BioImaging national research infrastructure, sample freezing conditions and cryo-EM preliminary acquisitions were performed at the AFMB microscopy platform and on the IMM imaging platform (Dr. Artemis Kosta). We thank the PACA Bioinfo platform for computing support.

## Funding

This project has received funding from the European Research Council (ERC) under the European Union’s Horizon 2020 research and innovation program (grant agreement No 832601). Proteomic experiments were partly supported by ProFI (ANR-10-INBS-08-01) and GRAL, a program from the Chemistry Biology Health (CBH) Graduate School of University Grenoble Alpes (ANR-17-EURE-0003). France-BioImaging national research infrastructure (ANR-10-INBS-04). CDC gratefully acknowledges STARPLUS 2020 (project no. 21CUNINACEPIGC042) from the University of Napoli for financial support.

## Data and materials availability

Genome sequences of the mutants of mimivirus reunion have been deposited to NCBI (accession numbers: KO_qu_946: OQ700912; KO_qu143: OQ700913, 2KO: OQ700914). 3D reconstruction maps and the corresponding PDB have been deposited to EMDB (Deposition number Cl1: 8ORS, EMD-17131; Cl2: 8ORH, EMD-17125). The mass spectrometry proteomics data have been deposited to the ProteomeXchange Consortium via the PRIDE (46) partner repository with the dataset identifier PXD041298. AlphaFold predictions were performed using HPC/AI resources from GENCI-IDRIS (Grant 2022-AD011013526).

## Authors contributions

C.A. conceived and designed the research; J-M.A., H.B., A.V., S.S., A.L., A.S., C.B. A.N., L.B., A.A., O.P. C.D.C., Y.C., and C.A. performed research; D.P. granted access to microscopy platform and helped optimizing sample freezing with A.V.; J-M. A and A.L. developed methodologies for genomic fiber and fibrils extraction and purification; H.B. developed the genetic tools and analyzed mutants; C.B. and A.L. built the Nqu143-GFP mutant and analyzed the data; A.V., collected CryoEM data; A.V., S.S., A.S., and C.A. analyzed CryoEM data Y.C., L.B., A.A., performed MS-based proteomics and analyzed the data; A.N. and C.D.C., performed GC-MS and NMR analyses and analyzed the data, H.B., A.V., A.S., S.S. A.N., C.D.C., Y.C.and C.A. wrote the manuscript.

## Competing interests

Authors declare that they have no competing interests

## Notes

### Competing Interest Statement

The authors have declared no competing interest.

### Summary of Updates

Revised version of the manuscript after review by two external reviewers. Figures have been modified, Title has changed and major rewriting of the abstract, introduction and discussion sections. More details have been provided in the result section. The two external reviewers were satisfied by this improved version.

