## Supplementary text and Figures for "Functional redundancy revealed by the deletion of the mimivirus GMC-oxidoreductase genes"

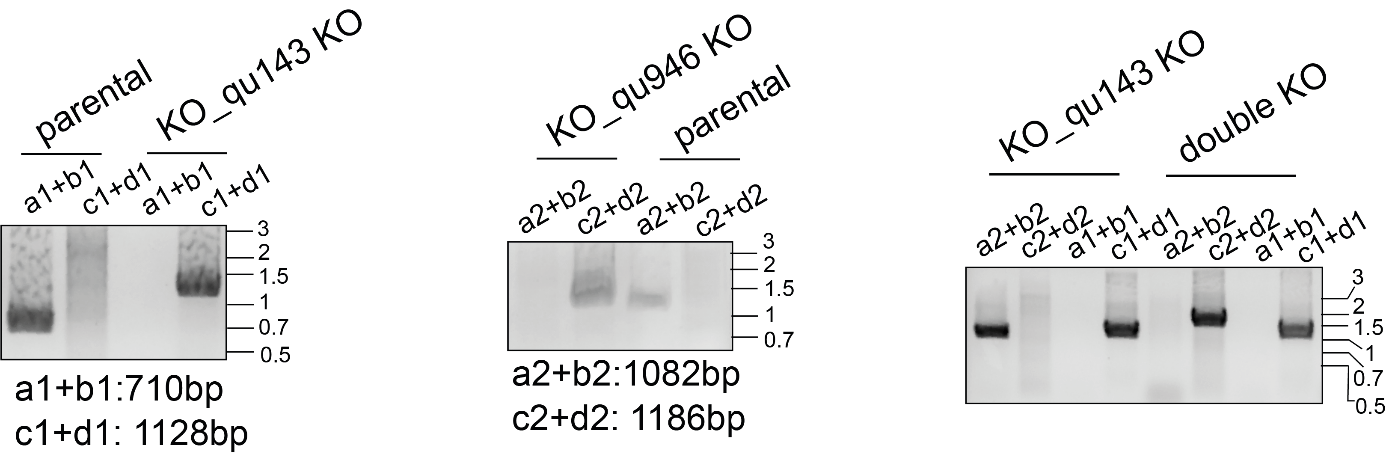

**Fig. S1: PCR of the genomic locus of qu_143 and qu_946** in single mutants and double KO demonstrates correct integration and clonality.

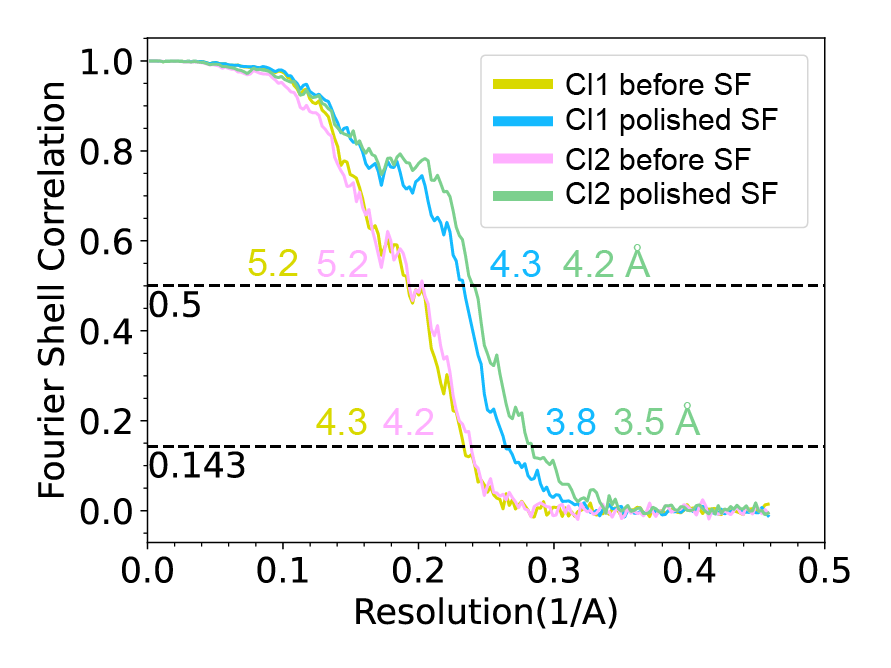

**Fig. S2: Fourier shell correlation (FSC) curves** for the final 3D reconstructions of the Cl1 and Cl2 helical assemblies.

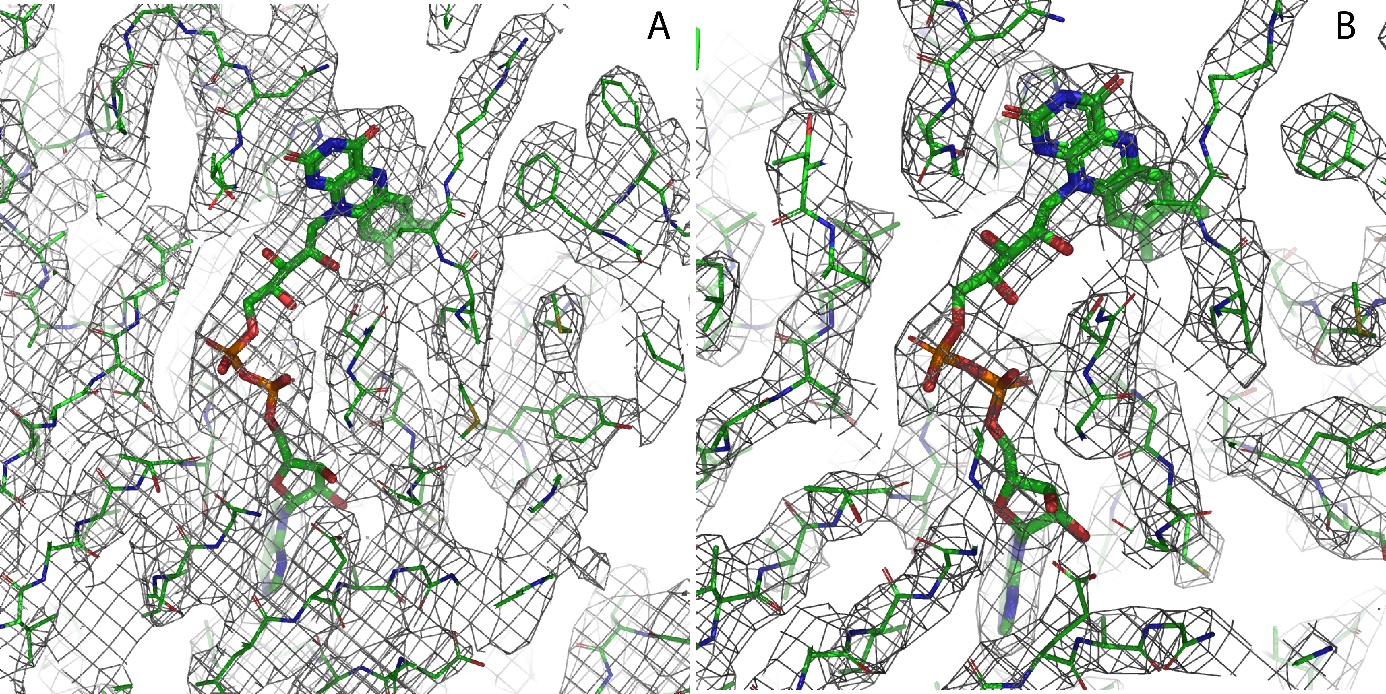

**Fig. S3:** **Zoom into the A] Cl1 and B] Cl2 maps** illustrating the fit of the side chains and the FAD ligand in the asymmetric unit.

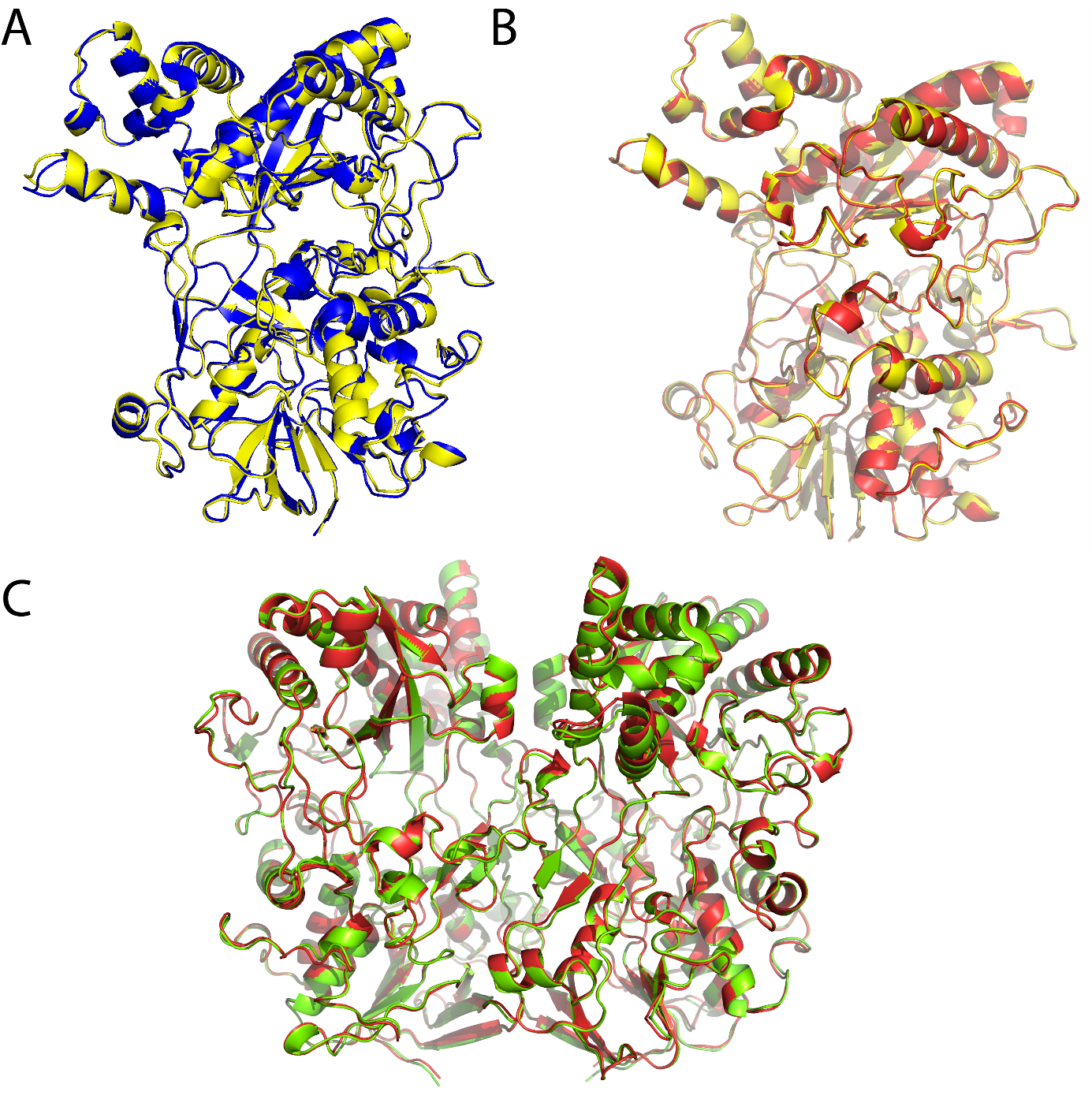

**Fig. S4: Structural superimposition of the qu_143 GMC-oxidoreductase of KO_qu946** A] Cl1 monomers (Chain A yellow, B blue), B] Cl2 monomers (chain A yellow, B red) and C] dimers of Cl1(yellow) and Cl2 (red) structures. Structural alignment was performed based on Cα superimposition using Pymol.

**A**

MKNRECCKCYNPCEKICVNYSTTDVAFERPNPCKPTPCKPTPIPCDPCHNTMVSKGEELFTGVVPILVELDGDVNGHKFSVSGEGEGDATYGKLTLKFICTTGKLPVPWPTLVTTLTYGVQCFSRYPDHMKQHDFFKSAMPEGYVQERTIFFKDDGNYKTRAEVKFEGDTLVNRIELKGIDFKEDGNILGHKLEYNYNSHNVYIMADKQKNGIKVNFKIRHNIEDGSVQLADHYQQNTPIGDGPVLLPDNHYLSTQSALSKDPNEKRDHMVLLEFVTAAGITLGMDELYK

**B**

MKNRECCKCYNPCEKICVNYSTTDVAFERPNPCKPTPCKPTPIPCDPCHNTMVSKGEELFTGVVPILVELDGDVNGHKFSVSGEGEGDATYGKLTLKFICTTGKLPVPWPTLVTTLTYGVQCFSRYPDHMKQHDFFKSAMPEGYVQERTIFFKDDGNYKTRAEVKFEGDTLVNRIELKGIDFKEDGNILGHKLEYNYNSHNVYIMADKQKNGIKVNFKIRHNIEDGSVQLADHYQQNTPIGDGPVLLPDNHYLSTQSALSKDPNEKRDHMVLLEFVTAAGITLGMDELYK

**Fig. S5: MS-based proteomic coverage of Nqu143-GFP in virions and fibrils.** Coverages obtained from analysis of A] purified virions and B] enriched fibrils. The residues covered by peptides identified in MS-based proteomic analyses are highlighted. The N-terminal part of qu_143 before the GFP sequence is underlined.

>qu_143 1^st^ in mimivirus reunion LP

MKNRE**CC**K**C**YN**PC**EKI**C**VNYSTTDVAFER**P**N**PC**K**P**T**PC**K**P**T**P**I**PC**D**PC**HNTKDNLTGDIV

IIGAGAAGSLLAHYLARFSNMKIILLEAGHSHFND**P**VVTD**P**MGFFGKYN**PP**NENISMSQN

**P**SYSWQGAQE**P**NTGAYGNR**P**IIAHGMGFGGSTMINRLNLVVGGRTVFDNDW**P**VGWKYDDV

KNYFRRVLVDIN**P**VRDNTKASITSVALDALRIIAEQQIASGE**P**VDFLLNKATGNV**P**NVEK

TT**P**DAV**P**LNLNDYEGVNSVVAFSSFYMGVNQLSDGNYIRKYAGNTYLNRNYVDENGRGIG

KFSGLRVVSDAVVDRIIFKGNRAVGVNYIDREGIMHYVKVNKEVVVTSGAFYT**P**TILQRS

GIGDFTYLSSIGVKNLVYNN**P**LVGTGLKNHYS**P**VTITRVHGE**P**SEVSRFLSNMAAN**P**TNM

GFKGLAELGFHRLD**P**NK**P**ANANTVTYRKYQLMMTAGVGI**P**AEQQYLSGLS**P**SSNNLFTLI

ADDIRFA**P**EGYIKIGT**P**NI**P**RDV**P**KIFFNTFVTYT**P**TSA**P**ADQQW**P**IAQKTLA**P**LISALL

GYDIIYQTLISMNQTARDSGFQVSLEMVY**P**LNDLIYKLHNGLATYGANWWHYFV**P**TLVGD

DT**P**AGREFADTLSKLSYY**P**RVGAHLDSHQG**C**S**C**SIGRTVDSNLKVIGTQNVRVADLSAAA

F**PP**GGNTWATASMIGARAVDLILGF**P**YLRDL**P**VNDV**P**ILNVN

>R135

MKNKE**CC**K**C**YN**PC**EKI**C**VNYSTTDVAFER**P**N**PC**K**P**I**PC**K**P**T**P**I**PC**D**PC**HNTKDNLTGDIV

IIGAGAAGSLLAHYLARFSNMKIILLEAGHSHFND**P**VVTD**P**MGFFGKYN**PP**NENISMSQN

**P**SYSWQGAQE**P**NTGAYGNR**P**IIAHGMGFGGSTMINRLNLVVGGRTVFDNDW**P**VGWKYDDV

KNYFRRVLVDIN**P**VRDNTKASITSVALDALRIIAEQQIASGE**P**VDFLLNKATGNV**P**NVEK

TT**P**DAV**P**LNLNDYEGVNSVVAFSSFYMGVNQLSDGNYIRKYAGNTYLNRNYVDENGRGIG

KFSGLRVVSDAVVDRIIFKGNRAVGVNYIDREGIMHYVKVNKEVVVTSGAFYT**P**TILQRS

GIGDFTYLSSIGVKNLVYNN**P**LVGTGLKNHYS**P**VTITRVHGE**P**SEVSRFLSNMAAN**P**TNM

GFKGLAELGFHRLD**P**NK**P**ANANTVTYRKYQLMMTAGVGI**P**AEQQYLSGLS**P**SSNNLFTLI

ADDIRFA**P**EGYIKIGT**P**NI**P**RDV**P**KIFFNTFVTYT**P**TSA**P**ADQQW**P**IAQKTLA**P**LISALL

GYDIIYQTLMSMNQTARDSGFQVSLEMVY**P**LNDLIYKLHNGLATYGANWWHYFV**P**TLVGD

DT**P**AGREFADTLSKLSYY**P**RVGAHLDSHQG**C**S**C**SIGRTVDSNLKVIGTQNVRVADLSAAA

F**PP**GGNTWATASMIGARAVDLILGF**P**YLRDL**P**VNDV**P**ILNVN

>Mg878

MTTEKKHISGKISGDIIIIGAGGAG**C**VLAHYLARFSRHKIVLIEAGSWHLND**P**VVSD**P**QG

FFGKYN**P**SNENVSMAQN**P**EYSWQA**P**LQ**P**DTGAYGKR**P**IIAHGMGVGGSTMVNRLNLVVGG

RTVFDQDW**P**ESWKYDNVKKYFQRILADIN**P**IRDNSEVNVTNSVLDALRILAQQQIDSGTY

VDFMLNKATGNV**P**NVEKTE**P**GSQLLNLND**P**EGINSVVAFSNFYFGVNKLADGTYIRKYGA

NTYLNNNYVDSNGHGIGEYSNLRVLAD**C**VVDRIIFKHRKAVGVNIIDINGNSHYVKAKQE

IVVSGGAFYT**P**TILQRSGIGDFDHLSKIGVQNLVYHNSLVGTGLKNHYS**P**ISSIKVNGTS

SEISKFLSNMTTT**P**ENMGFKGVAELGYHRLD**P**NK**P**SNANEVTYRKYQ**C**MVVNG**P**AI**P**SDQ

QFIYNISSSTGNYFGIITDDVRFA**P**EGYVKIAT**P**NI**P**RDA**P**KIFFNTFTNYTVTNE**P**AVD

QW**P**RAQKTLSALISAFLGYDIVYQIVQQMNIIVKNQGFNVDLEIAY**PP**NDLVSQLHDGLE

LYGENWWHYLV**P**GLALNNNSTNAEKQFADVLSKLSYF**P**RSGAHLDSHQG**C**T**C**AIGRVVDE

HLRVIGTENLRVVDLSAA**P**FL**P**GGNTWATASMIGARGVDIILKR**P**VLQSL**P**L**C**DV**P**KL**P**I

**C**NY

>mvi_825

MTTEKKHISGKISGDIIIIGAGGAG**C**VLAHYLARFSRHKIVLIEAGSWHLND**P**VVSD**P**QG

FFGKYN**P**SNENVSMAQN**P**EYSWQA**P**LQ**P**DTGAYGKR**P**IIAHGMGVGGSTMVNRLNLVVGG

RTVFDQDW**P**ESWKYDNVKKYFQRILADIN**P**IRDNSEVNVTNSVLDALRILAQQQIDSGTD

VDFMLNKATGNI**P**NVEKTE**P**GSQLLNLND**P**EGINSVVAFSNFYFGVNKLADGTYIRKYGA

NTYLNNNYVDSNGHGIGEYSNLRVLAD**C**VVDRIIFKHRKAVGVNIIDINGNSHYVKAKQE

IVVSGGAFYT**P**TILQRSGIGDFDHLSKIGVQNLVYHNSLVGTGLKNHYS**P**ISSIKVNGTS

SEISKFLSNMTTT**P**ENMGFKGVAELGYHRLD**P**NK**P**SNANEVTYRKYQ**C**MVVNG**P**AI**P**SDQ

QFIYNISSSTGNYFGIITDDVRFA**P**EGYVKIAT**P**NI**P**RDA**P**KIFFNTFTNYTVTNE**P**AVD

QW**P**RAQKTLSALISAFLGYDIVYQIVQQMNIIVKNQGFNVDLEIAY**PP**NDLVSQLHDGLE

LYGENWWHYLV**P**GLALNNNSTNAEKQFADVLSKLSYF**P**RSGAHLDSHQG**C**T**C**AIGRVVDE

HLRVIGTENLRVVDLSAA**P**FL**P**GGNTWATASMIGARGVDIILKR**P**VLQSL**P**L**C**DV**P**KL**P**I

**C**NY

>mm_715

M**C**ELKIT**C**SRKKLSADIVIAGAGGAG**C**ILAHYLARFSNYKIILVEAGSWHLQD**P**NVSD**P**Q

GFFGKYN**PP**NENIRMSQN**P**AYAWQA**P**LE**P**DTGSYGNRAIVAHGMGIGGSTMINQLNLVVG

GRTVFDQDW**P**TGWKYDDIKKYFQRVLADID**P**IRDTTRVNITETILDSLRILAEEQINSGV

**P**VDYLLNKATGDV**P**NIEKTD**P**DSN**P**LNLNDIEGINSVVGFSNFYFGVNKLEDGSYIRKYG

GNTYLNRNYVDKNGRGIG**P**YSNLRVLSNIIVDRILFDNNKATGINVIDINGNSFQIKAKR

EVIISGGTFYT**P**TILQRSGIGDFGYLSQVGVKKLVYHN**P**LVGTGLKNHYS**P**MSTIRVTGS

SEEVANFLSNMQTT**P**ENMGFKGVAELGYHRLD**P**NK**P**DNADQVTYRKYQ**C**MVVGG**P**GI**P**TD

QQVVFNLTSGNFFTIITDDVRFA**P**EGYVKIAT**P**NI**P**RDA**P**KIFFNTFTTYT**P**TSA**P**ANEQ

W**P**TAQKTLAALISSLLGYDIVYQIIGQMNKLVADKGFNVKLEMAY**PP**NDLVSKLHEGLNT

FGENWWHYFV**P**KLAISDSSSSEEIEFADTLSKLSYF**P**RSGAHLDSHQG**C**T**C**RIGQVVDKK

LKVIGTDNLRVVDLSVA**P**FL**P**GGNTWATASMIGAHAVDLILGR**P**VLKEL**P**LDDI**P**ELFQN

N

>ma_782

M**C**ELKIT**C**SRKKLSADIVIAGAGGAG**C**ILAHYLARFSNYKIILVEAGSWHLQD**P**NVSD**P**Q

GFFGKYN**PP**NENIRMAQN**P**SYAWQA**P**LE**P**DTGSYGKRSVVAHGMGIGGSTMINQLNLVVG

GRTVFDQDW**P**SGWKYDDIKRYFQRVLADIN**P**IRDTTRVNVTETILDSLRILAEEQINSGV

**P**VDYLLNKATGDI**P**NIEKTD**P**DSN**P**LNLNDIEGINSVVGFSNFYFGVNKLEDGSYIRKYG

GNTYLNKNYIDKNGRGIG**P**YSNLRVLSNTIVDRILFNDNKATGINVIDTSGNSFQIKAKR

EVIISGGTFYT**P**TILQRSGIGDFGYLSQVGVKELVYHN**P**LVGTGLKNHYS**P**MSTIRVTGS

SEEVANFLSNMQTT**P**ENMGFKGVAELGYHRLD**P**NK**P**DNADQVTYRKYQ**C**MVVGG**P**GI**P**TD

QQVVFNLTSGNFFTIITDDVRFA**P**EGYVKIAT**P**NI**P**RDA**P**KIFFNTFTTYT**P**TSA**P**ANEQ

W**P**IAQKTLGALISSLLGYDIVYQIISQMNKLVADKGFNVKLEMAY**PP**NDLVSKLHEGLNT

FGENWWHYFV**P**KLAISDSSSSEEIEFADTLSKLSYF**P**RSGAHLDSHQG**C**T**C**RIGQVVNKK

LKVIGTENLRVVDLSVA**P**FL**P**GGNTWATASMIGAHAVDLILGR**P**VLKEL**P**LDDV**P**ELFQN

N

>qu_465 1^st^ in KO_qu_143 N**P**

MRKLSWQHIVLIVLAIILILWIISLLL**C**RK**P**VR**P**TYQV**P**IIQ**P**MQVIQ**P**HQNDID**P**AWQT

TYS**P**NNTDNQNQQYVLYYFNN**P**S**CP**H**C**KNFSSTWDMLKNNFRSINNLSLKEISTDKQENE

HLVFYYNIRRV**P**TIILVT**P**DKNLEYSGNKSLEDLTQFIRSNMNQ

>R443

MRKLSWQHIVLIVLAIILILWIISLLL**C**RK**P**VR**P**TYQV**P**IIQ**P**MQVIQ**P**HQNDID**P**AWQT

TYS**P**NNTDNQNQQYVLYYFNN**P**S**CP**H**C**KNFSSTWDMLKNNFRSINNLSLKEISTDKQENE

HLVFYYNIRRV**P**TIILVT**P**DKNLEYSGNKSLEDLTQFIRSNMNQ

>mg446

MRNLSWQHILIIIIIIIIIFWLVSWLFF**P**KNINV**P**VG**P**YY**P**VQ**P**QLIQ**P**MTIIT**P**NLGQN

MTGNQL**P**RNNVVSTTN**P**VNTESQ**P**SNTQD**P**FMLYYFHA**P**S**C**VH**C**RNFN**P**AWEMLRERLAG

SRGISTAKVDATK**P**ENENLVFYYNVSAF**P**TIILIT**P**DQNVEYNGNRT**P**DDLHNFVVAHIN

EHNNRVSK

>mvi_415

MRNLSWQHILIIIIIIIIIFWLVSWLFF**P**KNINV**P**TGSYY**P**VQ**P**QLIQ**P**MTIIT**P**

NLGQNMTGNQL**P**RNNVV**P**TIN**P**VNTESQ**P**SNTQD**P**FMLYYFHA**P**S**C**VH**C**RNFN**P**A

WEMLRERLAGSRGISTAKVDATK**P**ENENLVFYYNVSAF**P**TIILIT**P**DQNVEYNGN

RT**P**DDLHNFVVAHINEHNNRVSK

>mm_337

MRNLSWQHIVLIIIIIIIVFWLISWLFF**P**KSV**P**GTTG**P**YYQ**P**QIIQ**P**MTIL**P**QNNGVA**P**Q

YSYGAV**P**NDNMQNTGNTDMQNSVND**P**FVLYYFHS**P**T**C**GH**C**KNFN**P**AWELLQQKLSGSGGV

SAKSIDTTK**P**ENENLAFYYNVSAV**P**TIILIT**P**DRNVEYSGNRS**P**DDLYNFVIAHLNDHNN

RNRS

>ma_373

MRNLSWQHIVLIIIIIIIVFWLISWLFF**P**KTI**P**GTTG**P**YYQ**P**QIIQ**P**MTIL**P**QNNGAA**P**Q

YSYGTV**P**NDNMQNTSNTDMQTAVND**P**FVLYYFHS**P**A**C**GH**C**KNFN**P**AWELLQQKLSSSGGV

SAKSIDTTK**P**ENENLAFYYNVSAV**P**TIILIT**P**DRNVEYSGNRS**P**DDLYNFVIAHLNDHNN

RNRS

>qu_757 NP

MSWHTGSNQDNKLF**P**KGKLSGSYA**P**LDIAFENS**P**AMNEFENRL**C**HNN**P**IISERSMS**P**AVS

ASYSN**P**EATS**C**G**C**MQTQTQ**P**QTQHQTQHL**P**QTHHTDAHDQQKLSGIFYNRTTDAQNQFSE

TIN**PPP**SYTVHNTDIRI**P**LNRQQQY**P**ANHLGSELLEGYNNVGTE**PC**MGFWEILLLIILIA

VLVYGIYWLYKSE

>R710 NP

MSWHTGSNQDNKLF**P**KGKLSGSYA**P**LDIAFENS**P**AMNEFENRL**C**HNN**P**IISERSMS**P**AVS

ASYSN**P**EATS**C**G**C**MQTQTQ**P**QHQTLSQHL**P**QTHHTDAHDQQKLSGIFYNRTTDAQNQFSE

TIN**PPP**SYTVHNTDIRI**P**LNRQQQY**P**ANHLGSELLEGYNNVGTE**PC**MGFWEILLLIILIA

VLVYGIYWLYKSEK

>mg261 LP

MS**C**NISRDNNMTMRRDSMASLDTAYDSSNLMKDFQNKISNGNIHMNQNTGNNINDSIANL

IDSNKL**P**YQSEEKIDMNVEKIVGEYYNRDHDIQKQLDRDIIGNQYTH**P**NVSTMQNGVEF**P**

LNRSQYYS**P**EMHGSELLEGYSNVGS**C**MD**P**WRLILLIILIAALIYGLYWLYTNNKTNNLDF

>mvi_244 LP

MS**C**NISRDNNMTMRRDSMASLDTAYDSSNLMKDFQNKISNGNIHMNQNTGNNINDSIANL

IDSNKL**P**YQSEEKIDMNVEKIVGEYYNRDHDIQKQLDRDIIGNQYTH**P**NVSTMQNGVEF**P**

LNRSQYYS**P**EMHGSELLEGYSNVGS**C**MD**P**WRLILLIILIAALIYGLYWLYTNNKTNNLDF

>mm_173 LP

MA**C**NI**P**GDSTSVRNTMATLDTAYNNSNLMRDFQNKLSGNNTLSNNINDNISQLRNNQIL**P**

HQSEEKIDVNVDKIVGEYYNRDDDIKRQLNRDSTGNEYIY**P**GV**P**TVQNGVNF**P**LDRSQYY

S**P**QVYGSELLEGYTNTGSGIDFWRLLLLIILIVALIYGIYWLYNNNKSSL**P**

>ma_211 LP

MA**C**NI**P**GDSTSVRNTMATLDTAYNNSNLMRDFQNKLSGNNTLSNNINDNISQLRNNQIL**P**

HQSEEKIDVNVDKIVGEYYNRDDDIKRQLNRDSTGNEYIY**P**NV**P**TAQNGVNF**P**LDRSQYY

S**P**QVYGSELLEGYTNTGSGIDFWRLLLLIILIVALIYGIYWLYNNNKSSL**P**

>qu_734 1^st^ in 2KO and M4 LP

MS**C**QNYQSYG**C**GTY**PC**VTYGN**C**YTT**C**AS**PC**L**P**Y**P**TN**C**VQV**C**TSSQ**PCP**S**PCP**I**P**V**PCP**VT

IVEYITTA**P**TATTIESS**P**TGMALT**P**I**P**VGSTSI**P**SGTVTVITGYAAT**P**VRSIGGITLNSA

LGQFTV**P**LAGSYLITGYIGFSYNAVGIREVYVYKVDGATSVITLISTDSRNTTATN**P**TYI

SYSAMDYFNAGDRIFIAAAQNSGSTITTTADNRIAITRMNRQ

>L688

MS**C**QNYQSYG**C**GTY**PC**VTYGN**C**YTT**C**AS**PC**L**P**Y**P**TN**C**VQV**C**TSSQ**PCP**S**PCP**I**P**V**PCP**VT

IVEYITTA**P**TATTIESS**P**TGMALT**P**I**P**VGSTSI**P**SGTVTVITGYAAT**P**VRSIGGITLNSA

LGQFTV**P**LAGSYLITGYIGFSYNAVGIREVYVYKVDGATSVITLISTDSRNTTATN**P**TYI

SYSAMDYFNAGDRIFIAAAQNSGSTITTTADNRIAITRMNRQ

>mg_239

MS**C**YNY**C**GY**P**ASY**C**Y**P**SYN**PC**QIS**C**I**P**Y**PP**V**C**Q**P**I**C**Q**P**V**C**Q**P**A**C**L**P**A**PCP**A**PCPP**A**P**IVV

EYSTTA**P**TATTI**P**SGTVGVA**P**T**P**I**P**AGSTVI**P**AGTVTVISGYSAV**P**TKNVGGITLNTTTN

QFTL**P**LAGRYLISSYIGISANSIGTRESYIYRVNGTTGVISLITSDSRNATAVG**P**TYITL

TTEDRFNAGDRIFFAVTQNSGSVLTTT**P**DNRFTITRL**C**N

>mvi_223

MS**C**YNY**C**GY**P**TSY**C**Y**P**SYN**PC**QIS**C**I**P**Y**PP**V**C**Q**P**V**C**Q**P**V**C**Q**P**A**C**L**P**A**PCP**A**PCPP**A**P**IVV

EYSTTA**P**TATTI**P**SGTVGVA**P**T**P**I**P**AGSTVI**P**AGTVTVISGYSAV**P**TKNVGGITLNTTTN

QFTL**P**LAGRYLISSYIGISANSIGTRESYIYRVNGTTGVISLITSDSRNATAVG**P**TYITL

TTEDRFNAGDRIFFAVTQNSGSVLTTT**P**DNRFTITRL**C**N

>mm_151

MA**C**NNL**C**GYA**C**Y**P**YY**C**ATYN**PC**QIA**C**V**P**Y**PP**V**C**QTA**C**I**P**SY**P**AI**CPP**T**PP**L**PPPP**IIVEY

ATTT**P**TGTTI**P**TGTVGV**PP**T**P**I**P**AGSTVI**P**AGTVTVISGYS**P**T**P**TRNIGGITLNTTTNQF

TL**P**LAGRYLITSFIGISAN**P**TGTRESYIYRVSGTTGVISLITTDSRNATDVG**P**TYINLAT

EDSFQAGDRIFFAVTQNSGTVLTTT**P**NSRFTITRLSS

>ma_191

MA**C**NNL**C**GYA**C**Y**P**YY**C**ATYN**PC**QIA**C**V**P**Y**PP**V**C**QTA**C**I**P**SY**P**AI**CPP**T**PP**L**PPPP**IIVEY

ATTT**P**TGTTI**P**TGTVGV**PP**T**P**I**P**AGSTVI**P**AGTVTVISGYS**P**T**P**TRNIGGITLNTTTNQF

TL**P**LAGRYLITSFIGISAN**P**TGTRESYIYRVSGTTGVISLITTDSRNATDVG**P**TYINLAT

EDSFQAGDRIFFAVTQNSGTVLTTT**P**NSRFTITRLS

>qu_482 1^st^ in Nqu143-GFP NP

MAGSFTRKMYDN**C**ATQQTTKQSTD**P**LELLLDVNKYVN**C**NNI**C**K**P**LAQRY**P**SSAQLVDVES

SLWGIDKLASR**C**DSSKH**P**F**C**AKNG**C**LLTND**P**RIA**P**HIT**P**YA**C**EWGHTGDNSVVTTNMKM**P**

SH**P**GYTL**P**N**P**NI**C**KDQTNGYYHNAVKS**P**NMTG**P**QHQTV**P**QHQVV**P**QHQSVIK**P**KADQLRY

QNIQNR**P**Y

>R459

MAGSFTRKMYDN**C**ATQQTTKQSTD**P**LELLLDVNKYVN**C**NNI**C**K**P**LAQRY**P**SSAQLVDVES

SLWGIDKLASR**C**DSSKH**P**F**C**AKNG**C**LLTND**P**RIA**P**HIT**P**YA**C**EWGHTGDNSVVTTNMKM**P**

SH**P**GYTL**P**N**P**NI**C**KDQTNGYYHNAVKS**P**NMTG**P**QHQVI**P**QHQVV**P**QHQTV**P**QHQSVIK**P**K

ADQLRYQNIQNR**P**Y

>mg428

MSGHFTRKMYDG**C**AAQQDIKQSTN**P**LELIMDVNKYVH**C**DNI**C**K**P**AREY**PP**NGALLVDVES

SLWGIDKLASR**C**DSAKH**P**F**C**G**P**NG**C**LLTND**P**RVRQHIT**P**YA**C**ERGKAGDNAVITTNMRM**P**

QH**P**GYTV**P**NANI**C**NTHNGYYSN**P**NSK**P**VYHSQQ**P**VTHSQQ**P**VYHSQQ**P**VTHSQQ**P**VTHSQ

Q**P**VYHSQQ**P**ITHSQQRIL**P**NIRNQQVVHSG**P**TVR

>mvi_397

MSGHFTRKMYDG**C**AAQQDIKQSTN**P**LELIMDVNKYVH**C**DNI**C**K**P**AREY**PP**NGALLVDVES

SLWGIDKLASR**C**DSAKH**P**F**C**G**P**NG**C**LLTND**P**RVRQHIT**P**YA**C**ERGKAGDNAVITTNMRM**P**

QH**P**GYTV**P**NTNI**C**NTHNGYYSN**P**NSK**P**VYHSQQ**P**VTHSQQ**P**VYHYQQ**P**ITHSQQ**P**VTHSQ

Q**P**VTHSQQ**P**VTHSQQ**P**ITHSQQ**P**ITHSQQRIL**P**NIRNQQVVHSG**P**AVR

>mm_321

MSGHFTRKMYDG**C**AAQQDLKQSTN**P**LELILDVNKYVH**C**DNI**C**K**P**AREY**PP**NGALLVDVES

SLWGIDKLASR**C**DSAKH**P**F**C**S**P**NG**C**LLTNDSRVA**P**HTT**P**YA**C**ERGHVGENAVVTTNMRM**P**

KH**P**GYTL**P**N**P**NI**C**STQNNGYYAN**P**NNRQNNQVLVN**P**NNRQNNQI**P**QNNQVLVNHALQNAV

AQNHALQNYGLQNHALQNQVAQNKIAQNNVVQNQSQNYLAQNQ**P**VLNY**P**QHRIL**P**NIRNQ

QA**P**VINAQN**C**GNL**P**VVR

>ma_357

MSGHFTRKMYDG**C**AAQQDLKQSTN**P**LELILDVNKYVH**C**DNI**C**K**P**AREY**PP**NGALLVDVES

SLWGIDKLASR**C**DSAKH**P**F**C**S**P**NG**C**LLTNDSRVA**P**HTT**P**YA**C**ERGHVGENAVVTTNMRM**P**

KH**P**GYTL**P**N**P**NI**C**STQNNGYYAN**P**NNRQNNQVLVN**P**SNRQNNQI**P**QNNQI**P**QNNQI**P**QNN

QVLVNHALQNTVAQNHALQNYGLQNHALQNQVAQNKIAQNNVIQNQSQNYLAQNQ**P**VLNY

**P**QHRIL**P**NIRNQQA**P**VINAQN**C**GNL**P**VVR

>qu_657 1^st^ in megavirus chilensis LP

MSVSRKRIDH**C**ND**C**ANKNGSGWRIVNVKDVTYRKER**C**SDVFKHE**P**R**C**Q**C**G**C**QDKHHDRHD

**PC**V**P**DE**C**SKTDKQLTIVSAVN**P**TSNLII**P**EAGVL**P**V**P**VNNVILNGWTLTT**P**DLLNSFN**P**S

TGVFTATESGDYEINLVLSFRSNASLSAATNLSNV**P**LVSIIDAATGL**P**LYEAVQGF**P**TTS

SLVDVEV**P**VI**PP**I**P**ISVTVTSVLGVGQVVLNLIVSLSAGQQVEILVSSNGLSHI**P**SSF**P**I

G**P**ATFTFNTGTSLIVKKVRNI**P**KVIYSL**C**

>R623

MSVSRKRIDH**C**ND**C**ANKNGSGWRIVNVKDVTYRKER**C**SDVFKHE**P**R**C**Q**C**G**C**QDKHHDRHD

**PC**V**P**DE**C**SKTDKQLTIVSAVN**P**TSNLII**P**EAGVL**P**V**P**VNNVILNGWTLTT**P**DLLNSFN**P**S

TGIFTATESGDYEINLVLSFKSSAFLNATENLSNV**P**QVSIIDAATGL**P**LNEAIQGF**P**TTS

SQVVVDV**P**VV**PP**I**P**VLVTVTSVLGVGQISLSIIISLSAGQQVEILVSSNGLSHI**P**SSF**P**V

G**P**ATFTFNTGTSLIVKKVRNI**P**KVIYSL**C**

>mg749

MSVRKR**C**VIND**C**SKN**C**GWKIVESDSEYVRSHKHHES**P**RSH**C**NDDHDH**C**HEVFD**PC**NRNNN

RNFQRTLISVIN**P**TNSILEI**P**IEFTNTFAARAIGVQQVIGNTVNISGWSDTL**P**DILDAFD

NTTGIYTA**P**ENGDYEFNLILNFKTSV**P**LTVNDAGTNI**P**IVEIYDVASGSTLAGGSILL**P**T

SNIQITI**PP**IASGEL**P**IEIEATNVLGSGQVVLTAII**P**LVAGQQVRVRANSNGMVYN**P**IQE

IIE**P**AFIDFN**P**NNSASRLTIQKVRNT**P**IIRYILN

>mvi_703

MSVRKR**C**VIND**C**SKN**C**GWKIVESDSEYVRSHKHHES**P**RSH**C**NDDHDH**C**HEVFD**PC**NRNNN

RNFQRTLISVIN**P**TNSILEI**P**IEFTNTFAARAIGVQQVIGNTVNISGWSDTL**P**DILDAFD

NTTGIYTA**P**ENGDYEFNLILNFKTSV**P**LTVNDAGTNI**P**IVEIYDVASGSTLAGGSILL**P**T

SNIQITI**PP**IASGEL**P**IEIEATNVLGSGQVVLTAII**P**LVAGQQVRVRANSNGMVYN**P**IQE

IIE**P**AFIDFN**P**NNSASRLTIQKVRNT**P**IIRYILN

>mm_599

MNVRRR**C**VNND**C**AKN**C**GWKIVESDNEYVRNRRRHDE**P**VR**PC**H**C**NDNDNEEQVFD**PC**NRNN

NRNLQRTLVSVTN**P**TGSILEI**P**VEFG**P**VTFAARANAQVNALQVIGNSVVLTGWSDAV**P**DI

LDAFDNITGTYTA**P**ESGDYEFELVANFRTSV**P**LTVNDLGTNI**P**ILEIFDVASGL**P**LVGGA

VNL**P**TATVQFTI**PP**ISSGEL**P**IEIETTSVLSNGQVVLTAIITLVAGQQVRVRANSNGLVY

N**P**LQEIVAS**P**AFINFN**P**VGA**P**TKLTIQKIRNT**P**IVRYIL

>ma_656

MNVRRR**C**VNND**C**AKN**C**GWKIVESDNEYVRNRRRHDE**P**VR**PC**H**C**NDNDNEEQVFD**PC**NRNN

NRNLQRTLVSVIN**P**TGSILEI**P**VEFG**P**VTFAARANAQVGALQVIGNSVVLTGWSDAV**P**DI

LDAFDNITGTYTA**P**ESGDYEFELVANFRTSV**P**LTVNDLGTNI**P**ILEIFDVASGL**P**LVGGA

VNL**P**TATVQFTI**PP**ISSGEL**P**IEIETTSVLSNGQVVLTAILTLVAGQQVRVRANSNGLTY

N**P**LQEIVAS**P**AFINFN**P**VGA**P**TKLTIQKIRNT**P**IVRYIL

>qu_600 1^st^ in megavirus vitis NP

MNHYDQYQKYKKKYLDLKNQLNNSSQYGGN**C**GNYGNNQFNNQFNSQATNRYQTGGVEFDL

KDEVAFWGRQMMEHLLLLHLGLDEEELKNSALQNHMDWKRYLTENFFSKGVN**P**G**P**DQAFL

TTNELEKIGLLNKNVVNGLIDQTIQYKSKLVKTLNSGQWVGWIY**P**AMAQHMLEEAEYFKR

KVNG**P**DYT**P**EQETKFVVHHHSTEMGATTQLLD**P**TEKDNIKIAKSYADI**C**MSKLSGRNKK**P**

F**P**KQWTSQEEAILRGQD**P**VDLATLMRISLKYSRELTQFAKETGQKIDSKQLKSIIH**P**VLA

HHIFREFYRFTKRLEQLGAQ

>L567

MNHYDQYQKYKKKYLDLKNQLNNSSQYGGN**C**GNYGNNQFNNQFNSQATNRYQTGGAEFDL

KDEVAFWGRQMMEHLLLLHLGLDEEELKNSALQNHMDWKRYLTENFFSKGVN**P**G**P**DQAFL

TTNELEKIGLLNKNVVNGLIDQTIQYKSKLVKTLNSGQWVGWIY**P**AMAQHMLEEAEYFKR

KVNG**P**DYT**P**EQETKFVVHHHSTEMGATTQLLD**P**TEKDNIKIAKSYADI**C**MSKLSGRNKK**P**

F**P**NQWTSQEEAILRGQD**P**VDLATLMRISLKYSRELTQFAKETGQKIDSKQLKSIIH**P**VLA

HHIFREFYRFTKRLEQLGAQ

>mg690 LP

MNNYEKYLKYKSKYLGLKNSNQYGGQRYILQDEIYFWSRQMMEHFLILHLGLDNDNLKNE

AFELHNKWKKFIDNNFTERGVK**P**DIDTVFLTQFDLAKLAEIDIDTVNRLIDATDKYKNKL

IDILEQGKWVGWIFVSMVEHMLKETMYFKRKING**P**DFDVQEEIHFINDHHATELAATAQM

ID**P**N**P**LQQKNIEIARSYAQKTMSLLKLSGS**P**IAIESSE**P**F**P**RQWN**P**RDEEILKGLE**P**SDQ

ATLLKISLKYSQELTDYAKDTGMKIDSGELKSIIH**P**LLAHHIYREIARMTNTLQQLSTQ

>mvi_646 LP

MNNYEKYLKYKSKYLGLKNSNQYGGQRYILQDEIYFWSRQMMEHFLILHLGLDNDNLKNG

AFELHNKWKKFIDNNFTERGVK**P**DIDTVFLTQFDLAKLAEIDIDTVNRLIDATDKYKNKL

IDILEQGKWVGWIFVSMVEHMLKETMYFKRKING**P**DFDVQEEIHFINDHHATELAATAQM

ID**P**N**P**LQQKNIEIARSYAQKTMSLLKLSGS**P**IAIESSE**P**F**P**RQWN**P**RDEEILKGLE**P**SDQ

ATLLKISLKYSQELTDYAKDTGMKIDSGELKSIIH**P**LLAHHIYREIARMTNTLQQLSTQ

>mm_555

MNNYDKYIKYKNKYLGLKKMQRGGQKY**P**VEDEIYFWSRQLMEHLMIIHLGLAEEQFSLKT

GLLRDRDLKKEAGDLQQKWKAFIDNNFGSKGIK**P**GLDQVFLTQEELAKLGDIDMNAVNRL

IDATDKYKSHLLNVLETGLWVGYIY**P**AMVEHMLQETLYFRRKLNG**P**AFS**P**QEEILYINNH

HGTEMAATAQMID**P**N**P**LQQRDIDIARAYANKTMSLLKLSGS**P**IAVESTA**P**F**P**RQWS**P**QDE

EILKGLQ**P**SDEATLLAISLKYSQELTDYAHATGVRIDSGELKSIIH**P**LLAHHEYREFARF

TNTLEKLAAQR

>ma_609

MNNYDKYIKYKNKYLGLKKMQRGGQKY**P**VEDEIYFWSRQLMEHLMIIHLGLAEEQFSLKT

GLIRDRDLKKEAGDLQQKWKAFIDNNFGSKGVK**P**GLDQVFLTQEELAKLGDIDMNAVNRL

IDATDKYKSHLLNVLETGLWVGYIY**P**AMVEHMLQETLYFRRKLNG**P**AFS**P**QEEILYINNH

HGTEMGATAQMID**P**N**P**LQQRDIDITRAYANKTMSLLKLSGS**P**IAVESTA**P**F**P**RQWN**P**QDE

EILKGLQ**P**SDEATLLAISLKYSQELTDYAHATGVRIDSGELKSIIH**P**LLAHHEYREFARF

TNTLEKLAAQR

qu_738 1^st^ in moumouvirus australiensis

MIKIYRMKFKKFVFKFIDHDKRNFTVV**C**VNVYANKATHEFAHDNDFSKKLEWKIKHFKHA

HALERRIHQLVKETYFRESTGSLDQFADFKSVKV**C**VKDKIVKINLGENQEGN**P**VYKQVKS

VSKHYHVFVRGTK**P**LNRREKGAYTHSMKVHDIHLTGNLDQGLEFAEL**C**NFSI**P**ESGIHSV

QSQSSVTQSLNGQNVN**P**GAVVTGGDNWLSATNNANWNSTANTNAAWNSMNRNSVAQNSAS

KNANNWNSAANSAVKSSQNNNLSAMNNSLYNNNKAVNTNTINSTNNRNVSSQNNANRNAS

MATTYNNSVNSANSINTANTRSQTGGQDEEDFEKKYKKYKNKYAKLKNQKTSNF

>R692

MIKIYRMKFKKFVFKFIDHDKRNFTVV**C**VNVYANKATHEFAHDNDFSKKLEWKIKHFKHA

HALERRIHQLVKETYFRESTGSLDQFADFKSVKV**C**VKDKIVKINLGENQEGN**P**VYKQVKS

VSKHYHVFVRGTK**P**LNRREKGAYTHSMKVHDIHLTGNLDQGLEFAEL**C**NFSI**P**ESGIHSV

QSQSSVTQSLNGQNVN**P**GAVVTGGDNWLSATNNANWNSTANTNAAWNSMNRNSVAQNSAS

KNANNWNSAANSAVKSSQNNNLSAMNNSLYNNNKAVNTNTINSTNNRNVSSQNNANRNAS

MATTYNNSVNSANSINTANTRSQTGGQDEEDFEKKYKKYKNKYAKLKNQKTSNF

>mg_243

MKKFVFKFVDHDKKNVTKINIGVFSDAKVEWTMDGVNDLQDHIQWEIHHIKRNHALNKEL

MRVIRQRYF**P**GVAESQNENQIGDYKIIKIKMKEDSTAVNINEESNQ**P**TFRQIKSFEM**C**VA

KYARGNK**P**TKHSDVHNYTKTHYVHEFSAGREVDQTINLTELATFIV**P**TNESKSTNNTVQN

RNNVTGGDNDDFMNNVRTSYAANRDRIYGSLDKN**P**GNNSTVYRNTFNADRNTNTMDNLQK

NNTNNNNKTNYNNHWNKIVSNRNSQTGGVHNDQEYEEKYKKYKEKYLQLKNQKAKNMRI

>mvi_227

MKKFVFKFVDHDKKNVTKINIGVFSDAKVEWTMDGVNDLQDHIQWEIHHIKRNHALNKEL

MRVIRQRYF**P**GVAESQNENQIGDYKIIKIKMKEDSTAVNINEESNQ**P**TFRQIKSFEM**C**VA

KYARGNK**P**TKHSDVHNYTKTHYVHEFSAGREVDQTINLTELATFIV**P**TNESKSTNNAVQN

RNNVTGGDNDDFMNNVRTSYAANRDRIYGSLDKN**P**GNNSTVYRNTFNADRNTNTMDNLQK

NNTNNNNKTNYNNHWNKIVSNRNSQTGGVHNDQEYEEKYKKYKEKYLQLKNQKAKNMRI

>mm_155

MKKYIFKFVDHERRAVTKISVGVFVNKKNEEH**P**N**P**DDLDAHIAHAIQHIKRNHILKRELM

RIVAERYFNNASSTNGRFGEYILKKHHHKKESDLVNISNDGSQ**P**RFRKVKEFEMHKVKFT

RGRK**P**AEIAQIFDYTKTHYVHEFTITNESDQGVQLAELASFTV**P**INNEARTAAVNATTNR

NQATV**P**GTLNATGGDDD**P**FMSVRQSYASNRGRMNSNNNNSYSSNNWDNNNTSTGSRSTAN

ANRNANTMSGNTSGNTSMYSANNNRNTLNGSSANANWNNVAASNRGSMANTSALNNNNNQ

MGGDNDDIYKDRYQKYKKKYLELKNQGLSNF

>ma_195 LP

MKKYIFKFVDHERRAVTKISVGVFVNKKNEEH**P**N**P**DDLDAHIAHAIQHIKRNHILKRELM

RIVAERYFNNASSTNGRFGEYILKKHHHKKESDLVNISNDASQ**P**RFRKVKEFEMHKVKFT

RGKK**P**AEIGQIFDYTKTHYVHEFAITNESDQGVQLAELASFTV**P**INNEARIVAANATTNR

NQATV**P**STLNATGGDDD**P**FMSVRQSYASNRSRTNNNNNSYSSNNWDNNNTSTGSRSTVNA

NRNANTMSGNTSGNTSMYSANNNRNTLNGSSANANWNNVAASNRSSIANTSALNNNNNQM

GGDNDDIYKDRYQKYKKKYLELKNQGLSNF

>mm_751 1^st^ in moumouvirus maliensis LP

MSYYNDFRDRDY**CC**GYISN**P**TITDRFYALDNNNNILSFSATRNNLGNYTQLGDSLR**C**H**P**I

SGLL**P**GQTAVGIAFR**P**ANRTLYLLVRTLTGGRLYTLNISDK**C**GAIAN**P**VGFGLITAGGA**P**

IVLTGTAFSISFN**P**TIDRLRVVSNTGQNLTVN**P**DNGVTIINTNLSYAVGDINFGNL**P**AVG

GIAYTNNYVGAGSTTLYDIATNQNSLVIQN**PP**NDGTLNTVGLLNIAVSQFLGFTIVNRSN

TAIAILRTGTQTGIYNINLLTGTATLLRRIV**P**DQ**C**NNGVIIGLAGV**P**SNRVL

>ma_819

MSYYNDFSYRDD**CC**GYISN**P**TITDRFYALDNNNNILSFSATRNNLGNYTQLGDSLR**C**H**P**I

SGLL**P**GQTAVGIAFR**P**ANRTLYLLVRTLTGGRLYTLNISEK**C**GAIAN**P**VGFGLITAGGA**P**

IVLTGTAFSISFN**P**TIDRLRVVSNTGQNLTVN**P**DNGVTIINTNLSYAVGDINFGNL**P**AIG

GIAYTNNFVGAGSTTLYDIATNQNSLVIQN**PP**NDGTLNTVGLLNIAVSQFLGFTIVNRSN

TAIAILRTGTQTGIYNINLLTGAATLLRRII**P**DQ**C**NNGVIIGLAGV**P**SNRVL

>mvi_86

MGGIIVFFLNNVYIGSIMSSRNFF**C**KHLAQN**CC**DYYNDDDIYY**P**IIKDNFYALDNYNNII

SFYV**P**ENEQYVL**C**QKNYSIKNR**P**IRGLISGQEAVGIDYR**P**ANNRLYLLARTQQIAQLYIL

DIND**PC**EVIAT**P**VGTNLATETGTIIQLNGTSFGVDFN**P**VVDRLRVVSNTGQNLRIN**P**TNG

ITIIDGNLSFA**P**GDINAGKI**P**AVGGAAYTNSFSGTTTTTLYDIDTNQNVLVIQN**PP**NDGT

LNTVGSLGVNVSEFVGFDIVGRNNTAYAILRVDNKTGLYNINLLTGKATLLKHIIASGSD

NSS**C**RLIGLTVL**P**INRIF

>mg94

MGGIIVFFLNNVYIGSIMSSRNFF**C**KHLAQN**CC**DYYNDDDIYY**P**IIKDNFYALDNYNNII

SFYV**P**ENEQYIL**C**QKNYSIKNR**P**IRGLISGQEAVGIDYR**P**ANNRLYLLARTQQIAQLYIL

DIND**PC**EVIAT**P**VGTNLVTETGTVIQLNGTSFGVDFN**P**VVDRLRVVSNTGQNLRIN**P**TNG

ITIIDGNLSFA**P**GDINAGKI**P**AIGGAAYTNSFSGTTTTTLYDIDTNQNVLVIQN**PP**NDGT

LNTVGSLGVNISEFVGFDIVGINNTAYAILRVDNKTGLYNINLLTGKATLLKHIIASGSD

NSG**C**RLIGLTVL**P**INRIF

**Fig. S6: Sequences of the proteins ranked first in the fibrils of at least one virus and orthologous sequence in other viruses.** Cys and Pro are highlighted in red and cyan, to ease the reading. NP means no promoter in 5’ of the gene and LP, late promoter.

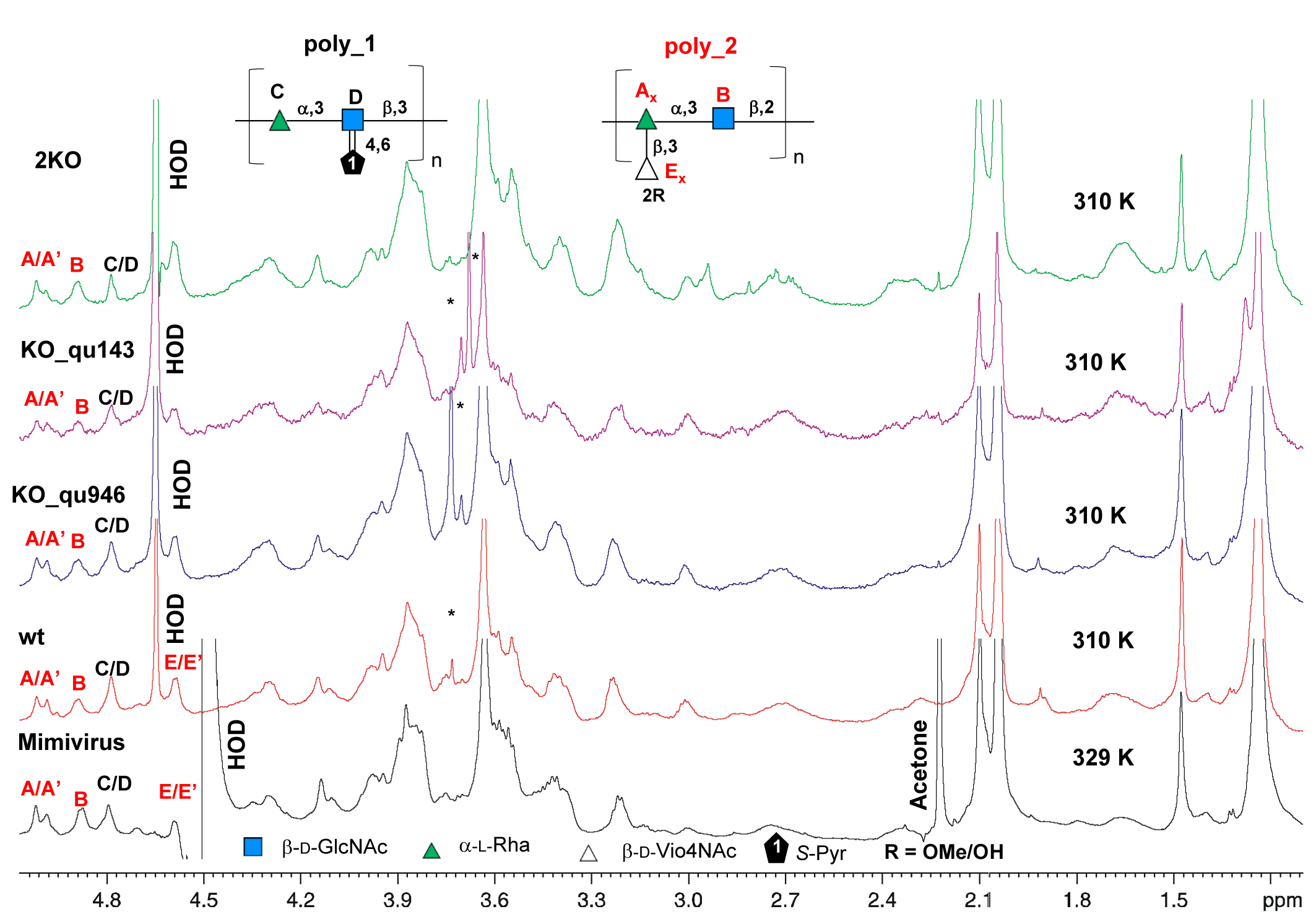
**Fig. S7: NMR analysis of the fibrils of mimivirus reunion wt and mutants compared to mimivirus.** The ^1^H NMR spectra were recorded on a Bruker 600 MHz instrument, in D_2_O, and the temperature is specified on the spectrum. The anomeric signals related to the poly_1 (C and D units) are shown in black, while those connected to the poly_2 are shown in red. Above the NMR spectra are shown the structures of the 2 polysaccharides as reported in Notaro *et al*, 2021. * indicates impurities related to traces of DTT left after the purification procedure.

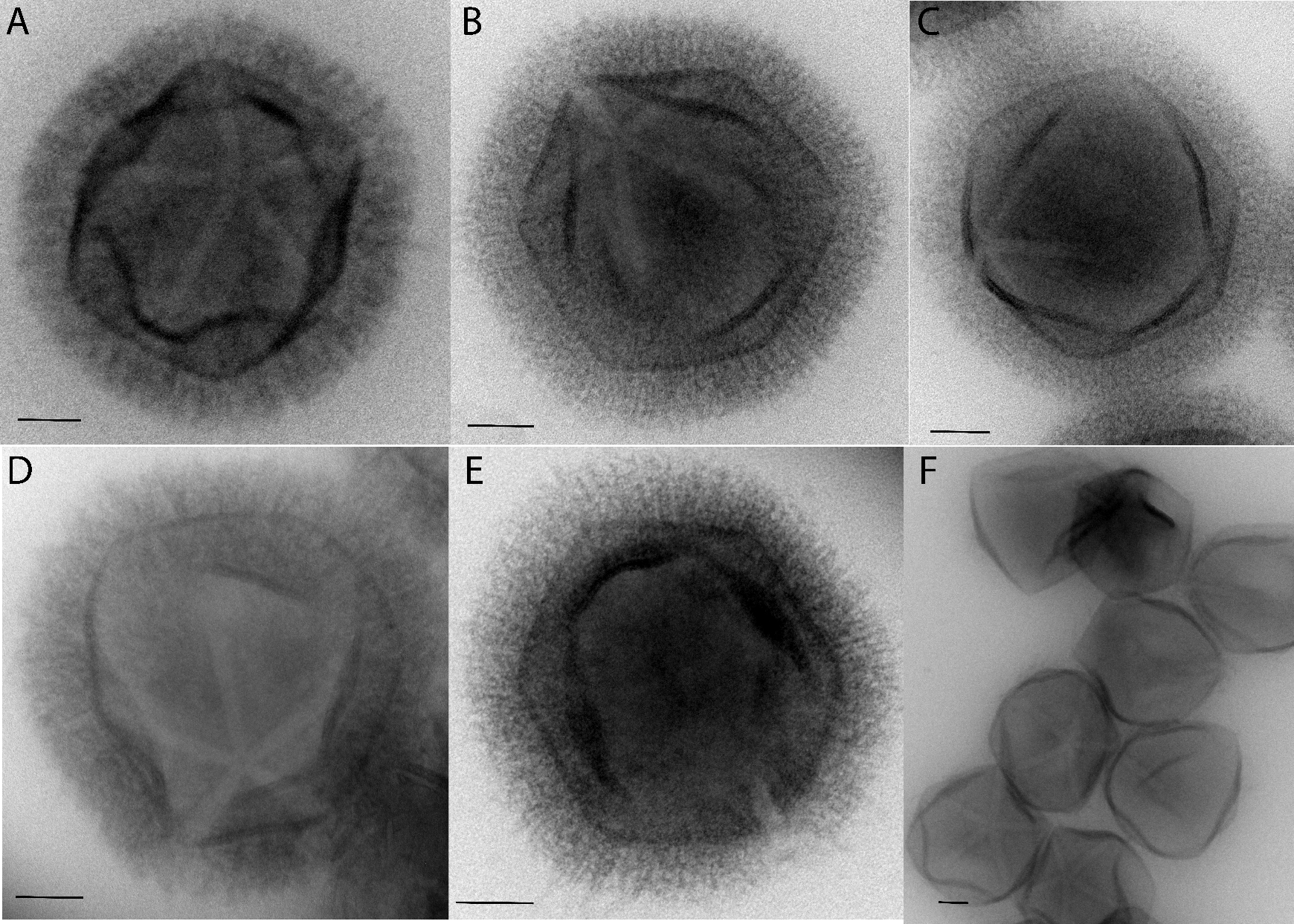

**Fig. S8: NS-TEM images** of A] mimivirus reunion wt, B] KO_qu946, C] KO_qu143, D] 2KO, E] Nqu143-GFP, F] M4 virions stained with uranyl acetate coupled with methylcellulose showing the same appearance of the reticulated layer of fibrils, with thicknesses ranging from 80 to 90 nm (scale bar 100 nm).

| **Table S1: List of primers used in this study.** | | |
| --- | --- | --- |
| HB161 | promoter mg153 | CTTTTGCAAAAAGCTTCGTCATCAAAATTAATAACTTG |
| HB162 | promoter mg153 | gtCAGAAGAATCAAGTTTAATTATTTAATGGTAACTAACT |
| HB163 | promoter mg741 | CTTTTGCAAAAAGCTTAAAATATTAGCAATTTTATCATAT |
| HB164 | promoter mg741 | gtCAGAAGAATCAAGTAAGTCGATATTCGATGT |
| HB168 | primer to introduce Not site + 3UTR megavirus | gcggccgcAGCGATAAGGATCCGCGT |
| HB169 | primer to introduce Not site + 3UTR megavirus | TTAGGGGCAGGGCATGCT |
| HB170 | primer to introduce Not site + 3UTR megavirus | ATGCCCTGCCCCTAATAATATTATTTTTAAATAATCAAAT |
| HB171 | primer to introduce Not site + 3UTR megavirus | TATCGCTgcggccgcTCTAAAGGACTATTTTGAA |
| HB190 | genotyping cassette integration KO | TCTACATGAGCATGCCCTGC |
| HB191 | KO q946 3 Homology arm | TCCTTTAGAGCGGCCGTTACGACAGGATCATTAAAATG |
| HB192 | KO q946 3 Homology arm | CTTATCGCTGCGGCCgcGGAATCTTCTTAACAAGTTATTTAG |
| HB193 | KO q946 5 Homology arm | CTTTTGCAAAAAGCTtGATCATTAGGAGGATACGCCATTTG |
| HB194 | KO q946 5 Homology arm | CTAATATTTTAAGCTGGTCCTGATGCTGCTGCA |
| HB195 | KO q143 3 Homology arm | TCCTTTAGAGCGGCCGGTTTAGCTACTTATGGAGCAAA |
| HB196 | KO q143 3 Homology arm | CTTATCGCTGCGGCCgcGTAATATTGATTTGACCGATAGACA |
| HB197 | KO q143 5 Homology arm | CTTTTGCAAAAAGCTtGTATAAAAACAGTGTCCCTAAATC |
| HB198 | KO q143 5 Homology arm | CTAATATTTTAAGCTCATTGGATCTGTTACAACA |
| HB233 | genotype q143 KO | GCCATCAAATTCCTTAACTCAC |
| HB234 | genotype q143 KO | GATTTATCTTCACCAGTATCTCTAG |
| HB235 | genotype q946 KO | TATGTGCGACAATATGTGCG |
| HB236 | genotype q946 KO | ATTGGTGTAATAGGTAACTGTTTGG |
| HB239 | genotype q143 KO | ATGCACCTGTATTAGGTTC |
| HB240 | genotype q946 KO | AGTTCTATTGGCGTACCG |
| CG 69 | qPCR nourseotricine | TACCACTCTTGACGACACGG |
| CG 70 | qPCR nourseotricine | AGTACGAGACGACCACGAAG |
| CG 71 | qPCR WT integration Queen | TCCTAAACCTCTTCAAGGAGAC |
| CG 72 | qPCR WT integration Queen | TACTGAACATTGACGCGAC |
|  | Add GFP to endogoenous tagging vector | CTTTTGCAAAAAGCTTATGGTGAGCAAGGGCGAGG |
|  | Add GFP to endogoenous tagging vector | attaaAAAATTATCATTACTTGTACAGCTCGTCCATGCC |
|  | Add N-terminal domanin q143 into GFP vector | gccctTGCTCACCATAGTATTATGACAGGGATCACATGGA |

**Table S3: KO_qu946 genomic fibers data statistics**

| **Mimivirus reunion KO_qu946 mutant** | **5-start fiber** | **6-start fiber** |
| --- | --- | --- |
|  | **Cl1** | **Cl2** |
| **Dimensions and symmetries** |  |  |
| Protein shell width  (external, internal) (Å) | 287.1-120.3 | 319.3-153.2 |
| Shell thickness (Å) | 84 | 83 |
| DNA ring diameter   (external, internal) (Å) | 132-93 | 163-124 |
| Spacing Shell-DNA | 5 | 5 |
| **3D Refinement** (Fig. 4) |  |  |
| Symmetry imposed | Helical | Helical, C3 |
| Helical parameters  (rise, twist) | 7.949 Å, 138.921° | 20.497 Å, 49.426° |
| Initial particle images (#) | 442,237 | 172,431 |
| Final particle images (#) | 98,882 | 20,899 |
| Initial model used | Featureless cylinder | Featureless cylinder |
| Model resolution   (Å) (masked) | 4.3 | 4.2 |
| FSC threshold | 0.5 | 0.5 |
| **Asymmetric unit model statistics** |  |  |
| *Model composition* |  |  |
| Non-hydrogen atoms | 101,24 | 101,24 |
| Protein residues | 1298 | 1298 |
| *R.m.s. deviations* |  |  |
| Bond lengths (Å) | 0.002 (0) | 0.003 (0) |
| Bond angles (°) | 0.506 (0) | 0.512 (0) |
| *Validation* |  |  |
| MolProbity score | 1.72 | 1.58 |
| Clashscore | 7.38 | 5.54 |
| Rotamers outliers (%) | 0 | 0 |
| *Ramachandran plot* |  |  |
| Favored (%) | 95.52 | 95.9 |
| Allowed (%) | 4.48 | 4.1 |
| Disallowed (%) | 0 | 0 |

**Table S4: RMSD between monomers and dimers in KO_qu946 Cl1and Cl2 helices and between KO_qu946 and wt helices**

| **RMSD between monomers (Å)** | | | | | | | | |
| --- | --- | --- | --- | --- | --- | --- | --- | --- |
|  | **Cl1_qu143 A** | **Cl1_qu143 B** | **Cl2_qu143 A** | **Cl2_qu143 B** | **Cl1a_wt A** | **Cl1a_wt B** | **Cl2_wt A** | **Cl2_wt B** |
| **Cl1_qu143 A** | 0 | 0.474 | 0.428 | 0.466 | 0.573 | 0.573 | 0.829 | 0.845 |
| **Cl1_qu143 B** |  | 0 | 0.44 | 0.413 | 0.538 | 0.538 | 0.801 | 0.826 |
| **Cl2_qu143 A** |  |  | 0 | 0.321 | 0.457 | 0.457 | 0.756 | 0.772 |
| **Cl2_qu143 B** |  |  |  | 0 | 0.43 | 0.43 | 0.706 | 0.726 |
| **Cl1a_wt A** |  |  |  |  | 0 | 0.001 | 0.78 | 0.791 |
| **Cl1a_wt B** |  |  |  |  |  | 0 | 0.78 | 0.791 |
| **Cl2_wt A** |  |  |  |  |  |  | 0 | 0.328 |
| **Cl2_wt B** |  |  |  |  |  |  |  | 0 |
| **RMSD between dimers (Å)** | | | | | | | | |
|  | **Cl1_qu143** | **Cl2_qu143** | **Cl1_wt** | **Cl2_wt** |  |  |  |  |
| **Cl1_qu143** | 0 | 0.467 | 1.912 | 0.704 |  |  |  |  |
| **Cl2_qu143** |  | 0 | 1.878 | 0.66 |  |  |  |  |
| **Cl1_wt** |  |  | 0 | 1.952 |  |  |  |  |

Cl1a_wt 7PTV; Cl2_wt 7yx3
